# Systematic exploration of predicted quaternary structures within pandemic-relevant viral proteomes

**DOI:** 10.64898/2026.09.29.755321

**Authors:** Yewon Han, Riya Narain, Razan Abbara, Marcelo Querino Lima Afonso, Jeannie Austin, Damian Bertoni, Sooyoung Cha, Alexander I. Cowen-Rivers, Joseph Ellaway, Davide Fransos, Kyle Gion, Darren Hsu, Chantal Hulo, Rachel Seongeun Kim, Oleg Kovalevskiy, Agata Laydon, Jack Mason, Sreenath Nair, Urmila Paramval, Nilkanth Patel, Risha Patel, Ivanna Pidruchna, Jeremy Ratcliff, Maxim Tsenkov, Niccolò Alberto Elia Venanzi, Melanie Vollmar, Newton Wahome, Elisa L.H. Wong, Jennifer Fleming, Sameer Velankar, Milot Mirdita, Philippe Le Mercier, Martin Steinegger, Christian Dallago, Joe Grove

## Abstract

Mechanistic understanding of viral protein-protein interactions enables global health security and pandemic preparedness, yet experimental characterisation remains difficult, costly, and often restricted to specialised laboratories. Here, we describe an *in silico* campaign using AlphaFold2 and AlphaFold-Multimer to predict monomers and dimer structures within 2,812 viral proteomes from 23 viral families relevant to human health. We report high-confidence predicted structures of 5,279 hetero-, and 2,749 homo-dimers. Structural clustering of interfaces reduces the set fivefold to 1,598 consolidated by form and function, of which 471 (29.5%) lacked detectable similarity to experimentally determined interfaces in the PDB. These clusters yielded two notable findings: new data on vaccinology-relevant glycoproteins, and accurate prediction of the specificity and cleavage-site recognition of viral proteases, which are targets for anti-viral development. Viral monomers and high-confidence dimers are available for interactive browsing in the Pandemic Preparedness Portal of the AlphaFold Database (https://alphafold.ebi.ac.uk/), and the full dataset is available for bulk download.

## Introduction

As exemplified by the COVID-19 pandemic (N. Zhu et al. 2020) viruses pose a persistent threat to global health and security. Mechanistic understanding of their biology can inform the development of interventions such as vaccines and therapeutics, mitigating pandemic risks (Wrapp et al. 2020). With small genomes, viruses rely on genomic economy and protein multi-functionality to ensure a high density of function despite a highly constrained coding capacity (Firth 2014). A common strategy for exerting and regulating function is multimeric self-assembly, driving the formation of both symmetrical structures - such as capsids, replication complexes, or glycoprotein spikes - and asymmetrical assemblies. Knowledge of the structure–function relationships of these complexes guides vaccinology, through rational immunogen design (McLellan et al. 2013) and structure-informed therapeutic development (Faysal et al. 2024).

Thus far, much of our understanding of protein complexes in virology has been provided through experimental structural biology and biochemical assays of direct interaction (Shah et al. 2022). The low throughput of these analyses has limited the scope of investigation to a relatively small number of tractable laboratory strains from priority viruses. Moreover, some viruses have proven recalcitrant to cell culture investigation (Kaur et al. 2026) or require biosafety controls that limit the scale of characterisation.

Existing knowledge of viral protein-protein interactions (PPIs) is distributed across several resources, though most focus on host–virus interactions rather than interactions between viral proteins themselves - for example, VirHostNet, the Host-Pathogen Interaction Database, and P-HIPSTer (Guirimand et al. 2015; Urban et al. 2025; Lasso et al. 2019). Resources describing intra-viral interactions are comparatively scarce, and include databases such as IntAct, BioGRID, the Protein Data Bank (PDB), Swiss-Prot and STRING, which provide some experimentally derived interaction data (Hermjakob et al. 2004; Stark et al. 2006; Szklarczyk et al. 2025; Burley et al. 2019; UniProt Consortium 2025). While these resources offer valuable snapshots of intra-viral interactions, they fall short of the systematic, proteome-wide coverage needed to reveal conserved interaction patterns across viral diversity - a gap that recent advances in structure prediction are positioned to fill.

The advent of highly accurate deep-learning protein structure prediction, for example AlphaFold2 and its multimeric extension, AlphaFold-Multimer (Jumper et al. 2021; Evans et al. 2021), has created the opportunity for *in silico* exploration of PPIs. Exhaustive investigations remain intractable for almost all cellular organisms due to the combinatorial explosion of pairwise searches across a large proteome. Consequently, previous studies have used strategies to reduce the search space; for example, applying classifiers to triage for likely PPIs, drawing on experimental data to select candidates, or pooling multiple protein sequences within each prediction (Humphreys et al. 2021; Han et al. 2026; Todor et al. 2026; J. Zhang et al. 2025; Schmid et al. 2025; Qi et al. 2026). Viruses, however, present an ideal testbed for systematic, at-scale investigation of PPIs, as their compact genomes are computationally tractable for proteome-level, all-vs-all searches. Indeed, this tractability was recently exploited to generate exhaustive AlphaFold-Multimer interactomes for human herpesviruses (T. Zhang et al. 2026; Soh et al. 2024). Extending this approach across a wide range of viral taxa would enable comparative analysis of protein complexes and their interaction modes.

In the present study we use an accelerated implementation AlphaFold-Multimer (Han et al. 2026; Didi et al. 2026) to perform prediction of intra-proteome PPIs across representative viral species from viral families known to infect humans. The rationale for this selection is that recent viral emergence has overwhelmingly occurred within viral families already known to include representatives capable of infecting humans (Woolhouse et al. 2012; J Woolhouse et al. 2013).

Accordingly, this dataset is designed to cover not only known human viruses, but also those with the potential to emerge in the human population. By sampling multiple representatives within each taxonomic group, our approach may uncover shared interactions within or between families or those that define particular genera, allowing explanations of virus biology and evolution based not only on the protein structures they possess (Mifsud et al. 2024) but also the interfaces these proteins create. Common interactions also present attractive targets for the development of broad-spectrum antivirals (Muratore et al. 2012; Link et al. 2020; Stray et al. 2005) .

We developed and tested a prediction pipeline against virus-specific benchmark sequences, and then embarked on a campaign of ∼1.7M pairwise predictions, across ∼42K protein sequences taken from 2,812 viral proteomes. Granular analysis of the frequency of high-confidence dimer predictions across our dataset suggests technical limitations and potential routes to overcoming these, and reveals patterns of protein interaction that correlate with underlying virus biology. Interface-based clustering consolidates our dataset into biologically coherent groups, permitting efficient exploration of form and function. We showcase clusters of vaccinology-relevant viral glycoprotein heterodimers, and complexes that accurately capture the interactions of viral proteases with native substrates. This dataset is made available via the Pandemic Preparedness Portal of the AlphaFold Database (AFDB) (Bertoni et al. 2026), providing new accessible knowledge for hypothesis generation and pandemic preparedness.

## Results

### An exhaustive proteome-level search of viral protein interactions

We selected 2,812 viral proteomes from 23 human-infecting viral families for intra-proteome all-vs.-all pairwise dimer predictions, yielding ∼1.7M complex structures containing ∼8K high-confidence models (**Fig. 1a-c; Methods**). In parallel, we also performed structure predictions for the ∼42K individual monomers in our sequence set; monomer confidence was highly concordant with Big Fantastic Virus Database (BFVD) v3 ((R. S. Kim et al. 2025; Litvin et al. 2025; Odai et al. 2025; R. S. Kim et al. 2026)) for shared non-polyprotein sequences (**Supp. Fig. 1**). Notably, this includes mature proteins from polyprotein-encoding viruses (**Supp. Fig. 2**) for which relatively few structures are available in the AFDB. More than 50K monomeric and high-confidence dimer models are accessible alongside existing viral structures (R. S. Kim et al. 2025; Litvin et al. 2025; Odai et al. 2025; R. S. Kim et al. 2026) via a new Pandemic Preparedness Portal accessible from the homepage of the AFDB (**Fig. 1d**).

**Figure 1:**
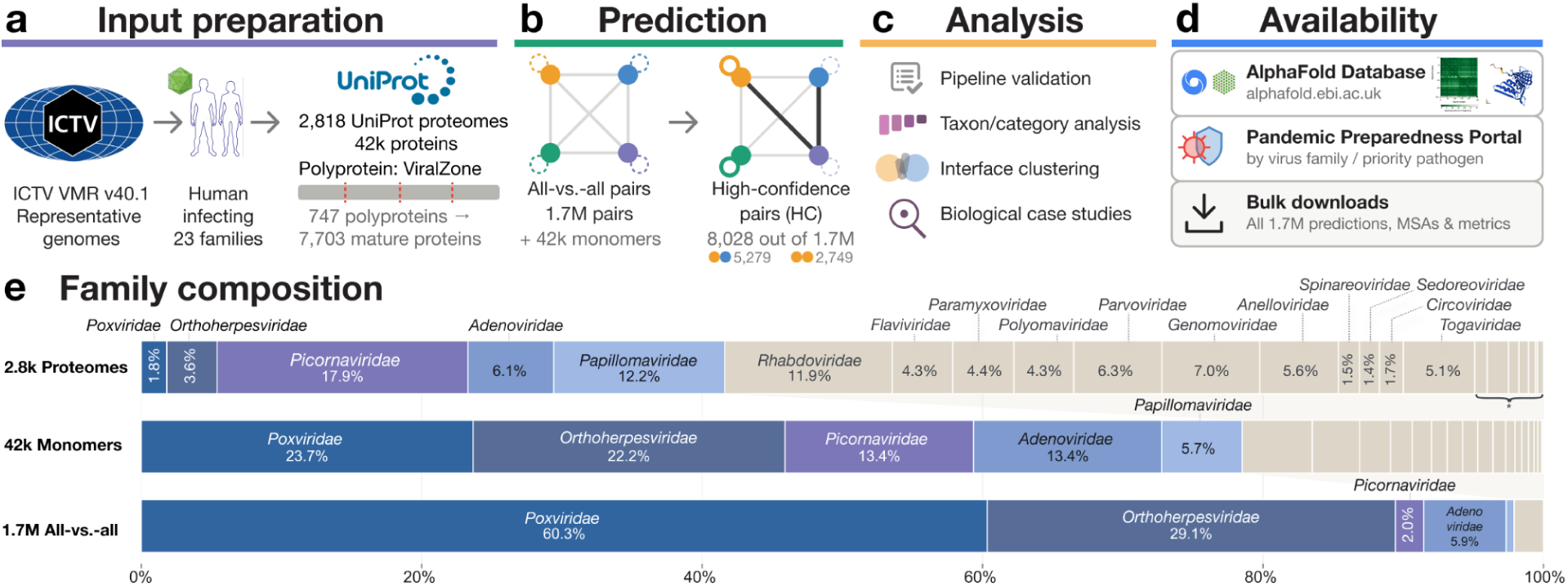
Proteome-wide dimer prediction for human-infecting viruses. **a**, We selected representative genomes of 23 human-infecting virus families from the International Committee on Taxonomy of Viruses Virus Metadata Resource (ICTV VMR) v40.1. **b**, We retrieved 2,818 Proteomes from UniProt and split polyproteins into mature peptides (ViralZone); all intra-proteome pairs were enumerated (1.7M pairs). **c**, The 1.7M dimeric and 42k monomeric structures were predicted (**Methods**), yielding 2,749 homodimers (6.75%) and 5,279 heterodimers (0.32%), together 8,028 dimers met *high-confidence (Han et al. 2026)* and were analysed for biological case studies. **d**, High-confidence dimers are available in the AlphaFold Database; all predictions via bulk download. **e**, Taxonomic breakdown of viral proteomes, monomeric proteins and all-vs-all pairs. Unlabelled segments (*) correspond, from left to right, to: Caliciviridae (0.9%), Hepadnaviridae (1.5%), Orthomyxoviridae (0.7%), Bornaviridae (0.7%) Filoviridae (0.5%), Pneumoviridae (0.1%) and Kolmioviridae (0.4%).

**Figure 2.**
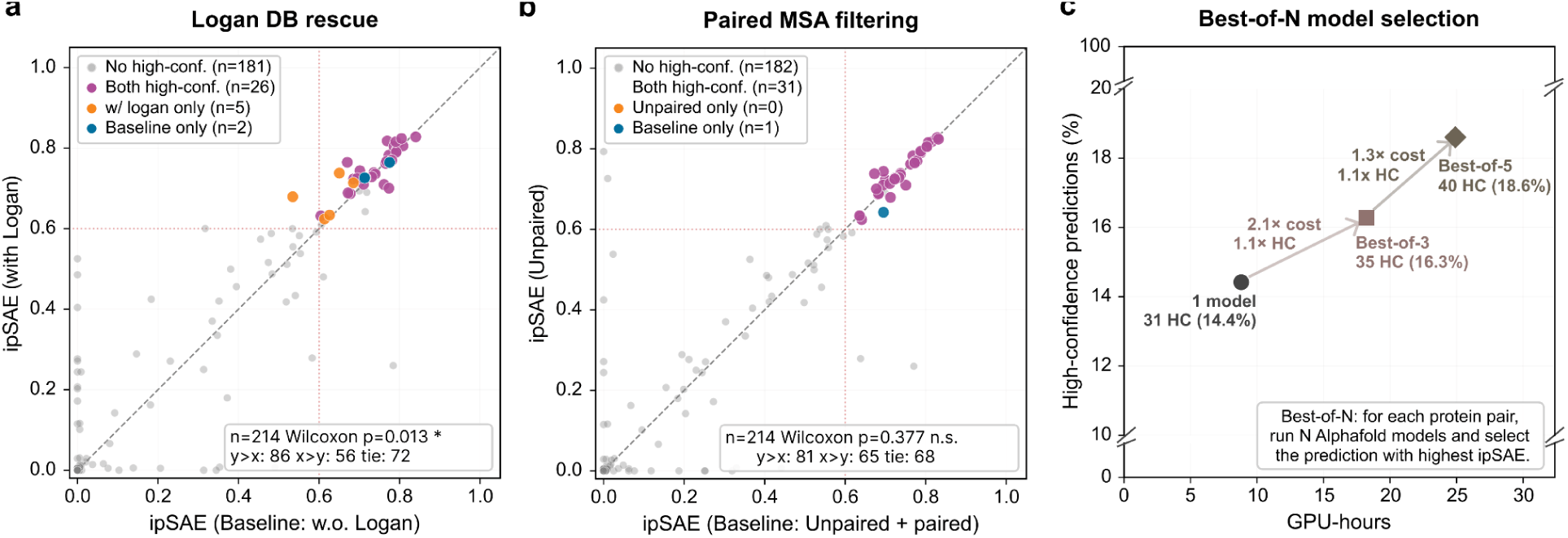
Effects of MSA composition and model selection on prediction yield and computational cost. Analyses used a set of 215 experimental viral complexes, with 214 evaluable pairs in a and b. High-confidence predictions satisfied ipSAE_max_ ≥ 0.60 and pDockQ2_max_ ≥ 0.23. In a and b: the production configuration is plotted on the y-axis; orange is high-confidence with the production configuration only, blue, with the alternative only, purple for both; dotted lines mark ipSAE = 0.6; P-values are from Wilcoxon signed-rank tests comparing paired ipSAE scores. **a,** Logan-block, enriched MSAs, concatenation improves prediction yield. ipSAE scores without (x-axis) and with (y-axis) the Logan block are compared (P = 0.013). Concatenation retained 26 high-confidence predictions, gained five and lost two. **b,** MSA pairing shows no significant change in ipSAE. Scores from paired-plus-unpaired MSAs (x-axis) are compared with those from unpaired-only MSAs (y-axis; P = 0.416). **c,** Best-of-*N* model selection offers modest gains but incurs increased computational cost. Using model 1 alone or selecting the highest-ipSAE prediction from three or five AlphaFold-Multimer models yielded 31, 35, and 40 high-confidence predictions at 1.0×, 2.1×, and 2.8× GPU cost, respectively.

The pool of viral species encompasses a wide diversity of genome compositions, coding strategies and proteome sizes; as a consequence of the combinatorial nature of our approach, DNA viruses with large genomes account for a high proportion of the pairwise predictions (**Fig. 1e**). For example, the *Poxviridae* contribute ∼2% of proteomes but ∼60% of all-vs-all pairs. The opposite holds for RNA viruses with small genomes: the *Picornaviridae* account for ∼18% of proteomes but ∼2% of all-vs-all pairs.

### Benchmark-based validation on the viral dimer prediction pipeline

Decisions about the prediction pipeline were underpinned by a systematic benchmarking against 215 experimental viral complexes deposited in PDB after the training of AlphaFold-Multimer and lacked dimeric structure-level similarity to the pre-training set (**Methods**; “*positive post-training benchmark set*”). Following Han et al. 2026, we considered a prediction successful if it had ipSAE_max_ ≥ 0.60 and pDockQ2_max_ ≥ 0.23 (Han et al. 2026; Dunbrack 2025; W. Zhu et al. 2023). For pipeline optimization, we compared prediction configurations by recall on the positive benchmark set at the established high-confidence thresholds, whereas the thresholds themselves were selected based on precision and evaluated separately using positive and negative benchmark pairs.

A common limitation when performing protein structure prediction across viral diversity is shallow multiple sequence alignments (MSAs) (Mifsud et al. 2024; Lytras et al. 2026). Recent work shows this can be mitigated through MSA enrichment using the Logan SRA-wide genome assembly (R. S. Kim et al. 2025; Chikhi et al. 2025). Pairwise concatenation of Logan-enriched MSA blocks (R. S. Kim et al. 2026) increased high-confidence yield from 28 to 31 out of 214, and improved ipSAE and pDockQ2 scores (Wilcoxon signed-rank test, P = 0.013; **Fig. 2a, Supp. Fig. 3a**). Meanwhile, adding filtered paired MSA to unpaired MSAs yielded 32 rather than 31 high-confidence predictions across 214 evaluable pairs, with no significant difference in ipSAE or pDockQ2 scores (Wilcoxon signed-rank test, P = 0.416; **Fig. 2b, Supp. Fig. 3b**).

**Figure 3.**
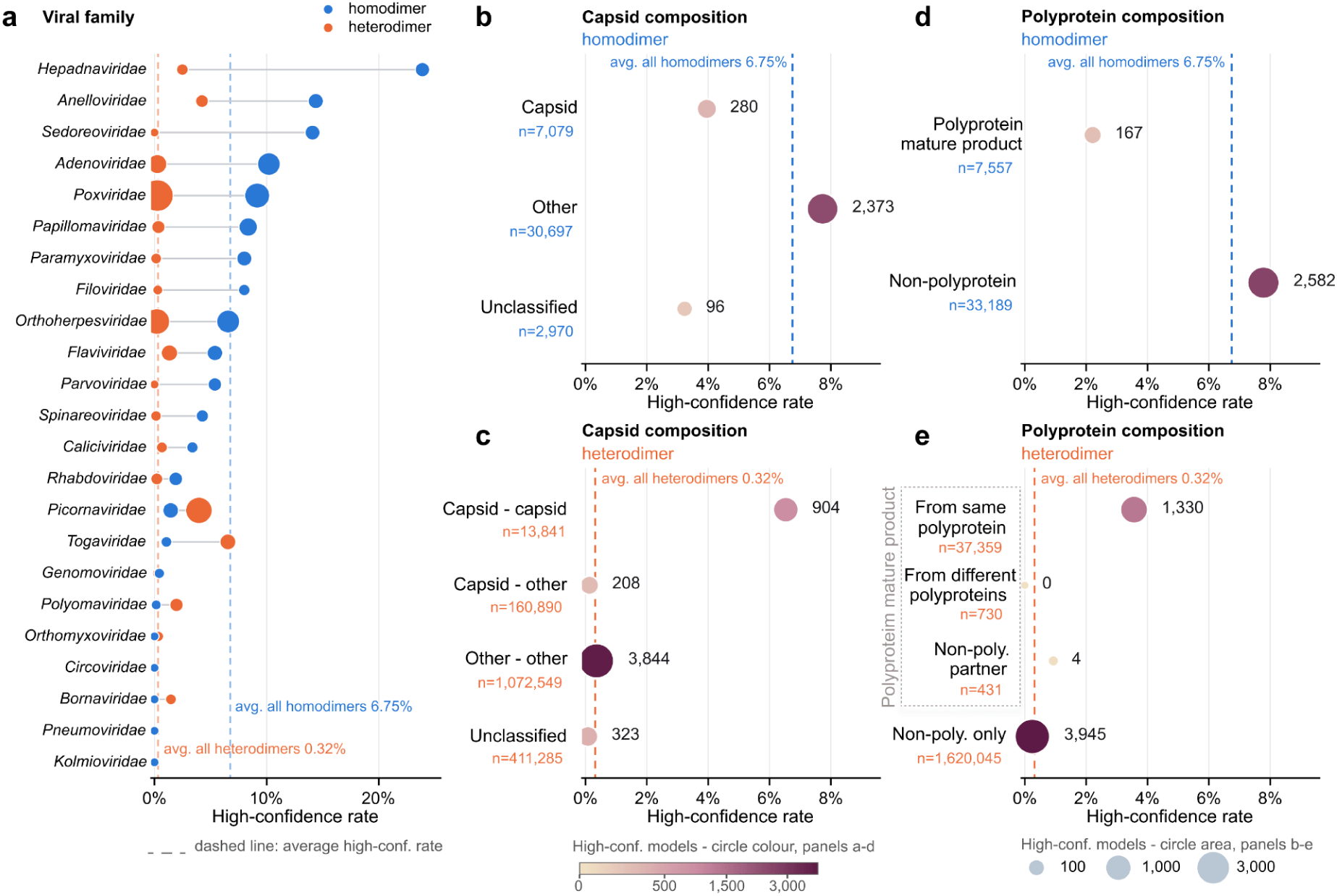
High-confidence fractions across viral dimer types, protein compositions and families. **a**, High-confidence fractions across all 23 NCBI taxonomic families, ordered by homodimer fraction. Connected dots compare dimer types within each family; colour identifies dimer type and circle area encodes high-confidence count as in b–e. In b-e, the dashed lines indicate overall homodimer and heterodimer fractions. **b,c,** High-confidence fractions by capsid composition for homodimers (b) and heterodimers (c), Other indicates non-capsid proteins; unclassified are entries with ambiguous protein identities. **d,e,** High-confidence fractions by polyprotein composition for homodimers (d) and heterodimers (e). In b–e, dot position indicates the high-confidence fraction; circle area and colour encode high-confidence counts using square-root scaling. Adjacent labels give high-confidence counts; n denotes input counts.

Beyond MSA configuration, we studied the effect of the number of AlphaFold-Multimer model weights sampled per prediction. Selecting the highest ipSAE prediction among one, three or five weight sets, resulted in 31, 35 and 40 high-confidence predictions, respectively. Runs with three or five weight sets added 2.1-fold or 2.8-fold GPU cost, and 1.1-fold or 1.3-fold high-confidence predictions compared to single-weight runs (**Fig. 2c**). High-confidence classification was also robust to recycling depth, with nearly unchanged yields across 4–20 recycles (**Supp. Fig. 4**).

Based on these results, we chose unpaired Logan-enriched MSAs and single-weight 12-recycle runs as our pipeline configuration. To evaluate the cutoff for the high-confidence predictions, we assembled a benchmark comprising positive post-training benchmark set together with structurally non-proximal experimental negative pairs (**Supp. Fig. 5a**). Applying the joint high-confidence criterion to the resulting predictions retained 31 of 215 experimentally supported positive pairs, and zero of 221 structurally non-proximal negative pairs, yielding no false-positives at a 14.4% recall (**Supp. Fig. 5b**). Recall falls below a recent AFDB complex benchmark (∼24%) (Han et al. 2026), which exclusively used dimeric assemblies, whereas in this study, we included dimeric chains extracted from larger heteromeric assemblies to increase sample size. This difference in benchmark composition, together with the inherent difficulty of structure prediction for some viral proteins (Lytras et al. 2026), may partly explain the lower recall.

### Patterns of high-confidence interactions across viral proteomes

We observe high-confidence fractions vary with dimer type and protein composition. Of 1,699,311 viral dimer predictions, 8,028 (0.47%) passed the joint confidence filter (**Fig. 1b**). Despite the recall rate (∼14%; Supp. **Fig. 5b**), this low high-confidence fraction reflects the exhaustive nature of the heterodimer campaign. We took an all-vs.-all approach in an attempt to uncover novel interacting proteins beyond annotations from existing databases, consequently heterodimers constituted the majority of the prediction candidates (97.6% of 1,699,311). Nonetheless, despite the vastly larger number of heterodimer candidates, homodimers constituted a substantial fraction of the high-confidence interactions recovered (34%; **Supp. Fig. 7**), consistent with the widespread use of homomeric self-association in viral protein complexes (Jayaraman et al. 2016).

In line with our exhaustive approach, heterodimer campaign yielded 0.32% high-confidence predictions, compared to 1% from our previous screen which filtered for dimers in STRING (Han et al. 2026). However, high-confidence yield was lower also for homodimer predictions (6.75% vs. 9.1% from our previous study (Han et al. 2026); **Supp. Fig. 6**), suggesting differences in protein composition or prediction difficulty for viral proteomes.

A greater high-confidence fraction for homodimers over heterodimers was observed across the majority of viral families with a few notable exceptions, for example, the *Toga-* and *Picornaviridae* (**Fig. 3a**). In these cases, the frequency of high-confidence heterodimers is increased by consistent prediction of glycoprotein assemblies (e.g. E1/E2, E2/E3) and components of the replication complex (nsp2/nsp4) in the *Togaviridae,* and capsid subunits (VP1/VP2, VP2/VP3) in the *Picornaviridae*.

Capsid proteins form contiguous protein shells and are a major structural component of non-enveloped viruses and many enveloped viruses. Capsid assembly is driven by repeating homo- and hetero-interactions that propagate across symmetry-constrained geometries, and is amenable to perturbation by anti-viral assembly inhibitors (Zlotnick and Mukhopadhyay 2011). Capsids, therefore, represent a paradigm of protein self-assembly - frequently forming both homo- and heteromeric configurations - and provide a useful reference point for our dataset. Unexpectedly, capsid homodimers had a lower high-confidence fraction than other–other (i.e. non-capsid) homodimers (3.96% versus 7.73%). By contrast, among heterodimers, capsid–capsid protein pairs showed an 18.2-fold greater high-confidence fraction than other–other pairs (6.53% versus 0.36%; **Fig. 3b,c**). This demonstrates that whilst our dataset recapitulates self-assembly of some capsid proteins, including diverse viruses that are poorly represented in the PDB **(Supp. Fig. 8)**, we are also likely to miss many capsid interactions, particularly homomeric ones. This may partly reflect the general difficulty of predicting protein-protein interactions, but capsids present an additional challenge: symmetry-related tiling means that the same capsid protein can participate in several geometrically distinct homomeric interfaces within the assembled shell (Rojas Labra et al. 2026). This is unlikely to be true for most non-capsid homomers in our dataset, and may create ambiguity when modelling simple dimers. Exploring higher-order stoichiometries may therefore rescue some of these missed homo-assemblies.

Polyprotein-derived homodimers similarly had a lower high-confidence fraction than non-polyprotein homodimers (2.21% versus 7.78%; **Fig. 3d**). As with capsids, this may partly reflect the failure to capture the relevant higher-order stoichiometries in assemblies commonly found in polyprotein-encoding viruses: capsids, glycoprotein spikes and lattices, and replication complexes, for example (den Boon et al. 2024; Haas et al. 2025).

Although many viruses (e.g., picornaviruses) encode a single polyprotein, some encode multiple distinct polyproteins across separate open reading frames or genome segments. We therefore asked whether high-confidence dimer frequency differed within and between polyproteins encoded by the same virus. Heterodimers comprising distinct mature products of the same polyprotein (i.e., intra-polyprotein interactions) reached 3.56% (1,330/37,359) high confidence, compared with no pairs (0/730) from inter-polyprotein interactions (**Fig. 3e**). The baseline for heterodimers not derived from polyproteins was 0.24%. This enrichment is consistent with polyproteins genetically consolidating functionally related proteins that evolved to act in concert. One clear example from our dataset is the structural polyprotein of the *Togaviridae,* which contains the self-assembling elements of both the capsid and the glycoprotein complex that together form the virion (Chen et al. 2018); we obtained 52 high-confidence complexes of structural proteins from 34 diverse togaviruses.

### Interface-clustering identifies functionally coherent groups of complexes

Searching for interaction interfaces that are conserved across viruses provides a complementary perspective on our dataset, grouping complexes by mode of interaction, rather than purely by composition or genetic relatedness. Following the approach of Strom et al. (Strom et al. 2026), we applied a stepwise clustering strategy (Methods); Foldseek-Multimer (W. Kim et al. 2025) clustering reduced 8,028 high-confidence complexes to 4,951 clusters (a 1.6-fold reduction), while subsequent Foldseek-Interface (Strom et al. 2026) clustering of their representatives yielded a final set of 1,598 interface clusters, a further 3.1-fold reduction and a 5.0-fold reduction overall (**Fig. 4a**).

**Figure 4.**
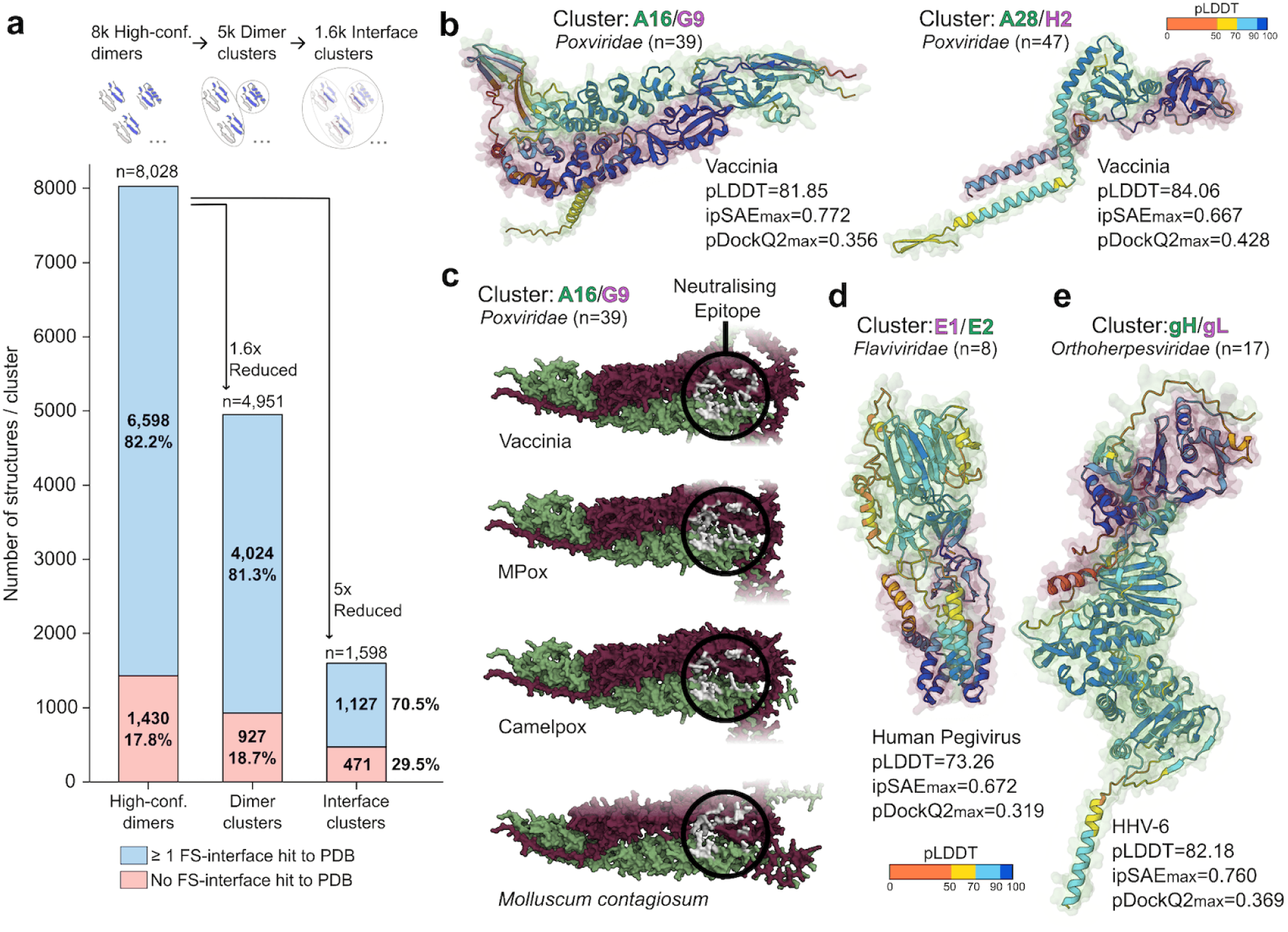
Clustering of the predicted viral interface space and glycoprotein complex example clusters. **a**, Foldseek-Multimer and Foldseek-Interface clustering of 8,028 high-confidence viral dimers (ipSAE_max_ ≥ 0.6; pDockQ2_max_ ≥ 0.23). Foldseek-Multimer clustering reduces the dataset 1.6-fold to 4,951 clusters, while Foldseek-Interface clustering of representatives yields a further 3.1-fold reduction to 1,598 clusters. Coverage of interfaces with a Foldseek-interface hit against PDB experimental interfaces is largely retained after Foldseek-Multimer clustering (82.2% to 81.3%); 471 Foldseek-Interface clusters (29.5%) have no PDB hit. **b,** Vaccinia heterodimers taken from A16/G9 (AF-0000000213241223) and A28/H2 (AF-0000000213229510) interface-clusters. Ribbon diagrams are colour-coded by pLDDT prediction confidence, as denoted in the key, with transparent surface representations highlighting individual chains colour-coded by protein identity, as shown in the complex labels. Annotations provide average pLDDT values and interaction confidence metrics. Cluster labels indicate complex identity, viral taxonomy and, in parentheses, number of members per cluster. **c,** Surface representations of A16/G9 dimers from diverse poxviruses (top to bottom: AF-0000000213241223, AF-0000000213009765, AF-0000000213150491, AF-0000000212886881) . A structurally conserved quaternary neutralizing epitope is highlighted in light grey. Surfaces are colour-coded by protein identity as indicated **d,e,** Ribbon diagrams of glycoprotein heteromers from interface-clusters of E1/E2 (*Flaviviridae*; AF-0000000212089925) and gH/gL (*Orthoherpesviridae*). HHV-6: human herpesvirus 6 (AF-0000000212432428).

This effectively consolidated complexes with shared interfaces, often reflecting a common function irrespective of sequence similarity or even protein composition. For example, the largest cluster (n=391) contained capsid VP2/VP3 heteromers from across the diverse *Picornaviridae*. Foldseek-Multimer initially separated these complexes into 95 clusters, largely correlated with viral genera, whereas interface clustering consolidated them into a single group.

Interfaces conserved across multiple viruses are natural candidates for core, functionally important contacts. Moreover, the structural diversity within individual clusters provides additional insight into how these interactions vary across related viruses. Most were conserved at the family level, consistent with viruses within a taxonomic group sharing fundamental features of virion architecture and replication. Such conserved interfaces may therefore represent attractive targets for broad-spectrum countermeasures. Only seven clusters contained similar dimers spanning multiple viral families. These were predominantly homodimers, from DNA viruses with cellular homologs, consistent with acquisition through horizontal gene transfer. For example, one dUTPase homodimer cluster (n=22) contained proteins of shared origins in cellular hosts (Baldo and McClure 1999; J. Li et al. 2024), from the *Adenoviridae*, *Orthoherpesviridae* and *Poxviridae* (**Supp. Fig. 9**).

### Clusters of vaccinology-relevant glycoprotein heterodimers

Viral structural proteins assemble into higher-order complexes that form virus particles, enabling their release from infected cells, transport to new host cells, and delivery of the viral genome upon entry (Prasad and Schmid 2012). These proteins frequently act through homo- or heteromeric assemblies and are major targets for adaptive immunity. Structural understanding of such complexes has underpinned rational design of vaccines to maximise protective responses, exemplified by the introduction of stabilising proline substitutions into class-I fusion proteins from RSV, SARS-CoV-2 and HIV (Sanders and Moore 2021). Accurate prediction of the quaternary organisation of viral structural proteins could therefore accelerate structure-informed vaccinology (McCool et al. 2026). As described above, the largest interface-clusters in our dataset comprise heteromeric complexes of *Picornaviridae* capsid proteins. However, these structures are already well represented in the PDB. To assess whether our dataset could instead recover structurally under-characterised viral surface complexes, we turned to viral entry glycoproteins.

Whilst many glycoproteins act as homotrimers (which are absent from our current analyses), our approach captured mechanistically relevant heterodimers, including components of the poxvirus membrane fusion machinery. The *Poxviridae*, particularly Mpox virus, pose an ongoing public-health threat, and have prompted renewed interest in vaccine development (Yu et al. 2025). Poxviruses possess a large entry fusion complex (EFC), containing at least 11 proteins, whose structure and mechanism of action remain incompletely understood. However, recent structural and immunological characterisation of the EFC provides valuable context for assessing our predictions (Lin et al. 2026; Meola et al. 2025; Yang et al. 2023).

Two large interface clusters contain the A16/G9 (n=39) and A28/H2 (n=47) heterodimers, which occupy the central and adaptor layers of the EFC, respectively (**Fig. 4b**) (Lin et al. 2026). Comparison with experimental structures deposited after the AlphaFold-Multimer training cut-off demonstrates accurate prediction of the heterodimeric arrangement of these mechanistically important proteins (**Supp. Fig. 10**).

A16/G9 is also an important target of the immune response: immunisation with the complex elicits protective responses, and monoclonal antibodies targeting it are neutralising—including antibodies that recognise quaternary epitopes formed at A16/G9 interface (Meola et al. 2025). One such epitope, identified in vaccinia virus, is structurally conserved across poxvirus A16/G9 dimers in our dataset (**Fig. 4c**), including highly divergent homologues such as those from *Molluscum contagiosum* (∼40% sequence similarity to vaccinia).

Beyond poxviruses, our high-confidence models provide structures for several glycoprotein heterodimers that are absent or poorly represented in the PDB. These include human pegivirus E1/E2 (**Fig. 4d**), which may exert a novel and currently uncharacterised membrane fusion mechanism (Mifsud et al. 2024), and human herpesvirus-6 gH/gL (**Fig. 4e**), which mediates interactions with host receptors (Tang et al. 2013). Together, these examples demonstrate how systematic structure prediction can recover conserved viral surface-protein assemblies of direct relevance to structure-informed vaccinology.

### Accurate prediction of viral protease-substrate engagement

Beyond grouping dimers by shared composition, interface-clusters can identify compositionally diverse groups that share a functional interaction mode. Two large, distinct clusters centred on viral cysteine proteases provide a striking example. Picornavirus 3C is a chymotrypsin-like protease (Seipelt et al. 1999), whereas adenovirus protease (L3/23k, AVP) has a unique fold (McGrath et al. 2003). In each interface-cluster, the protease was engaged with a diverse range of native viral substrates, which were consolidated by their shared mode of interaction rather than by protein composition (**Fig. 5**).

**Figure 5.**
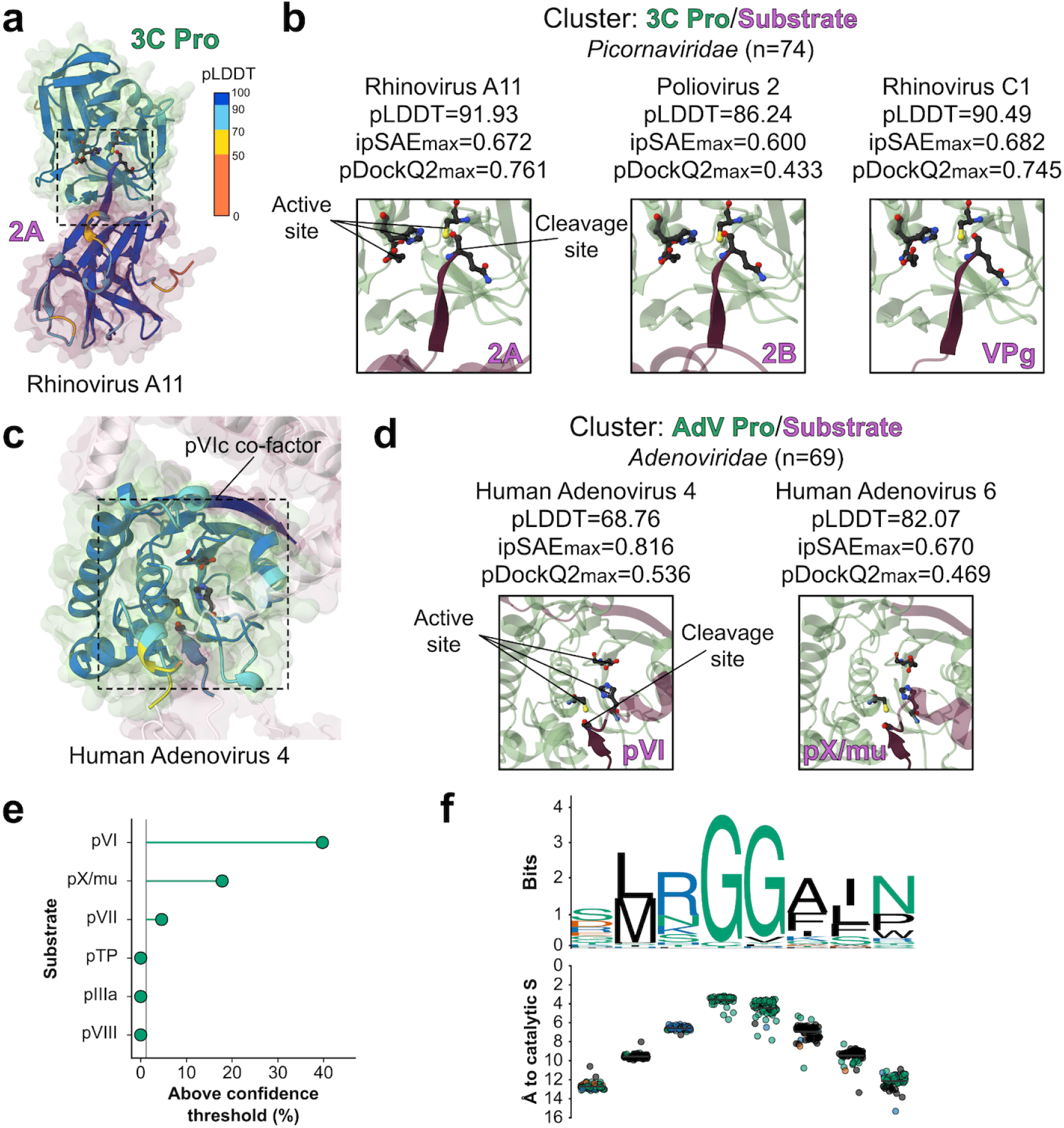
Modelling enzymatic activity of viral proteases. Highlighted residues in chain:number(s) notation. **a**, Ribbon diagram of the rhinovirus A11 3C protease in complex with its substrate 2A (AF-0000000212056371; A:145/B:40,71,147), coloured and annotated as in **Fig. 4b**; the dashed box indicates the area zoomed in the subsequent panel. **b**, Zoomed views of the 3C protease active site engaged with the substrate cleavage site, for rhinovirus A11 (substrate = 2A; A:141-145/B:40,71,147), poliovirus 2 (substrate = 2B; AF-0000000212064153; A:93-97/B:40,71,147) and rhinovirus C1 (substrate = VPg; AF-0000000212064278; A:18-22/B:40,71,147). In the zoomed views, ribbons show chain colour-coding only; the catalytic triad and the substrate P1 residue are drawn as sticks with heteroatom colouring, and everything beyond the cleavage site is shown in transparency. Cluster labels as in **Fig. 4**. **c**, Ribbon diagram of the human adenovirus (AdV) 4 protease in complex with its substrate pVI (AF-0000000212158930; A:29-33,230-242/B:56,73,123). The C-terminus of pVI (pVIc) is docked against the protease, recapitulating co-factor interactions. For clarity, low-pLDDT regions of pVI that do not contact the protease are faded to white. **d**, Zoomed views of the AdV protease active site, for human AdV 4 (substrate = pVI; A:29-34/B:56,73,123) and human AdV 6 (substrate = pX/mu; AF-0000000212195305; A:54,71,122/B:47-52), displayed as in b. **e**, Proportion of AdV protease/substrate predictions crossing our confidence thresholds for six classical substrates; the grey line indicates the baseline frequency for non-substrate proteins (1.2%). f, Sequence logo (upper) and matched per-residue distance from the catalytic sulfur (lower), for structurally aligned substrates in complex with the AdV protease (n=65 substrate/active site pairs).

Many viruses encode dedicated proteases that process viral proteins and polyproteins during replication and maturation, and can additionally target host proteins to manipulate cellular responses (X.-D. Li et al. 2005). These enzymes span several established protease folds and virus-specific architectures and display distinct substrate specificities (Rawlings and Bateman 2019). Their essential roles in replication have made viral proteases important targets for direct-acting antivirals, exemplified by therapies for HIV and hepatitis C virus (Bartenschlager et al. 2013).

Picornavirus 3C protease is principally responsible for processing the viral polyprotein and, because we used mature polyprotein products in our predictions, our dataset contains no intact 3C target sequences. Nonetheless, we obtained an interface-cluster of 3C from diverse members of the *Picornaviridae* engaged with three distinct cleavage products: VPg, 2A, and 2B. In each case, the C-terminal region of the cleavage product adopts a similar β-strand conformation at the entrance to the protease active site, positioning its terminal residue - the residue immediately preceding the cleaved peptide bond - adjacent to the catalytic cysteine (**Fig. 5a,b**). Therefore, these models effectively recreate the configuration of the protease and its product immediately following proteolytic cleavage.

AVP processes structural precursor proteins during adenovirus particle maturation. Unlike picornavirus 3C, complete native AVP substrates were present in our proteome-level pairings, and interface clustering grouped AVP complexes containing three of these substrates: pVI, pX/Mu and pVII. As observed for 3C, AVP substrates formed an upstream β-strand that positioned the cleavage site adjacent to the catalytic cysteine (**Fig. 5c,d**), recapitulating the geometry of proteolysis. Notably, pVI also serves as an AVP cofactor through its C-terminal peptide, pVIc, which activates the protease by remodelling the active site (Baniecki et al. 2013); this distinct interaction was also captured in our AVP/pVI models (**Fig. 5c**). Albeit the AVP-pVIc co-factor complex, in the absence of peptide substrate, was in the PDB at the time of AlphaFold-Multimer training (Ding et al. 1996).

To test whether these observations reflected authentic substrate recognition, we quantified AVP selectivity across the complete dataset, using the many non-substrate pairings as an internal negative control. Only ∼1% of AVP interactions with non-substrate proteins reached high confidence, compared with ∼40%, ∼20% and ∼5% for the native substrates pVI, pX/Mu and pVII, respectively (**Fig. 5e**). Thus, AVP showed substantial discrimination between substrates and non-substrates, although two additional native substrates, pIIIa and pVIII, yielded no high-confidence interactions. Viral proteases must recognise not only the correct substrate proteins but also specific cleavage sites within them. We therefore asked whether our predictions reproduced AVP sequence specificity by aligning substrate sequences according to their three-dimensional proximity to the catalytic site and generating sequence-logo plots alongside matched distance measurements (**Fig. 5f**). This recovered the key sequence features of a canonical AVP cleavage motif: (M/I/L)XGG–X. Taken together, these data demonstrate accurate and precise modelling of viral protease-substrate interactions.

Previous small-scale investigations have demonstrated that AlphaFold-Multimer can recapitulate interactions of cellular proteases with peptide targets (Zlobin and Golovin 2022; Veizaj et al. 2026; Ge et al. 2024). Here, we extend this principle to unbiased proteome-scale searches across diverse viruses, recovering substrate selection, cleavage-site specificity and mechanistically coherent interfaces. More broadly, the ability to systematically identify conserved molecular mechanisms across viral diversity may provide starting points for the discovery and design of antiviral inhibitors targeting shared viral vulnerabilities.

## Discussion

Building on other community-led efforts (R. S. Kim et al. 2025, 2026; Litvin et al. 2025; Mifsud et al. 2024; Odai et al. 2025; Nomburg et al. 2024; Soh et al. 2024), our study represents an exhaustive search of protein-protein interactions across diverse viral proteomes through AlphaFold-Multimer protein-structure prediction. While the wider virosphere remains a formidable target for systematic investigation, with ∼18K viral genomes taxonomically defined by the ICTV, our focus on viral families known to contain human-infecting viruses maximises the potential for hypothesis generation relevant to the development of anti-viral countermeasures. Moreover, computational analyses permit investigation of viruses irrespective of their amenability to growth in the laboratory, challenges of biocontainment, or availability of model systems and tools, thereby enabling exploration across broad viral diversity and potentially revealing targets for broad-spectrum intervention.

Guided by an experimentally grounded benchmark dataset, we employed a prediction pipeline that balanced accuracy, computational efficiency and the scale required for brute-force all-vs.-all searches. Stringent filtering (Han et al. 2026) yielded a high-confidence dataset with a low expected false-positive rate, but consequently a low overall frequency of retained models. While this is unsurprising given the combinatorial scale of our approach, patterns within the dataset also point to specific failure modes. For example: capsid proteins produced fewer high-confidence homomeric predictions than non-capsid proteins, despite their well-established propensity for self-assembly. One explanation is inadequate sampling of oligomeric space, such that some homomeric interactions may only be recovered at higher stoichiometries. The same may apply to viral glycoproteins, many of which form trimers. Exploring protein stoichiometry is therefore likely to be important for accurately modelling the symmetric assemblies that comprise virus particles and are particularly relevant to vaccine development. However, it should be noted that recent work suggests that AlphaFold may itself have limitations in modelling higher-order symmetric assemblies of viral structural proteins (Rojas Labra et al. 2026).

Deploying interface-based clustering (Strom et al. 2026) was highly effective in consolidating functionally related complexes across substantial sequence and structural divergence. This revealed conserved interaction modes shared across viral families and, in the case of proteases, mechanistically related assemblies formed by compositionally distinct proteins. We expect this approach to be particularly useful for navigating large and structurally diverse datasets and demonstrate its utility here by grouping highly diverse poxvirus glycoprotein assemblies.

Recent studies suggest that integrative structure prediction modelling of viral glycoproteins can illuminate interactions with the human immune response (Paciello et al. 2025) and guide immunogen design (McCool et al. 2026). Systematic prediction of glycoprotein complexes, together with other structurally important viral assemblies such as capsids and secreted proteins (Akey et al. 2014), could therefore provide a valuable stockpile of molecular knowledge for both proactive and outbreak-responsive vaccinology.

One capability exposed by our systematic approach is the ability of AlphaFold-Multimer to accurately model the substrate selectivity, sequence specificity and molecular mechanism of cysteine proteases from the *Picornaviridae* and *Adenoviridae*. Understanding whether this capability has arisen *de novo* or represents generalisation from distantly related proteases in the PDB, will require further investigation. Nonetheless, it could have broad applications, from investigating the molecular determinants of substrate specificity in diverse viruses to identifying previously unknown cleavage sites within viral proteins. Given the central role of proteolytic processing in the replication of many viruses, such mechanistic insight could also complement ongoing efforts to develop protease inhibitors as broad-spectrum antivirals (Nelen et al. 2026; Lithgo et al. 2024).

More widely, systematic interrogation of this and future datasets may reveal further unanticipated examples in which AlphaFold-Multimer captures molecular mechanisms or discovers novel interactions that have eluded comparable experimental screens. Our findings illustrate the potential for *in silico* exploration of protein-protein interactions in any viral proteome, revealing and consolidating knowledge of viral mechanisms, while offering a scalable route to rapidly analyse and respond to emerging pathogens.

The AlphaFoldDB Pandemic Preparedness Portal, through which predictions from this and future campaigns will be made available, represents a foundation for systematic exploration of structure, function and mechanism across diverse viruses. This has the potential to enable and accelerate fundamental discovery in molecular virology and, critically, the development of antiviral countermeasures: diagnostics, therapeutics and vaccines.

## Biosecurity statement

A general concern about virus research - whether *in vitro* or *in silico* - is that the practice may lead to manipulations of viruses that result in accidental or intentional increases in transmission or virulence. Cognisant of this, we limited our analysis strictly to interactions within each viral proteome, and chose not to explore interactions across proteomes, or between viral proteins and host proteins. Therefore, we may learn about how viral proteins assemble and, through this, how we might target viruses with drugs or vaccines. We will not learn about interactions between viral and host proteins that drive pathogenicity. The complexes are based on sequence data observed in nature, and we do not explore the impact of non-observed mutations on complex confidence. By making these choices, we aimed to maximise the potential benefits of our work, whilst minimising risks. Given this, the biosecurity risk associated with our dataset is low.

## Availability

High-confidence dimer, and all monomer predictions are available as individual entry pages through AlphaFold Database at alphafold.ebi.ac.uk. Dimers not passing the high-confidence threshold are provided at ftp.ebi.ac.uk/pub/databases/alphafold/collaborations/nvda/. The folding pipeline is available at github.com/NVIDIA-BioNeMo/BioNeMo-Structure-Prediction-Pipeline.

## Acknowledgements

We would like to thank NVIDIA colleagues, in particular Kyle Gion and Anthony Costa for support. EMBL-EBI also acknowledges their colleagues Oana Stroe, Gemma Wood, and Victoria Hatch for their support. We thank Lindsay Willmore and Anna Koivuniemi from Google DeepMind for their review and comments on the manuscript, as well as Evie Gray, Danielle Breen, and Myriam Khan for their support on communications. Martin Steinegger acknowledges support by the National Research Foundation of Korea (NRF) grants funded by the Korea government (MSIT) (RS-2024-00396026, RS-2025-00438101, RS-2026-25549295) and Novo Nordisk Foundation (NNF24SA0092560). Milot Mirdita acknowledges support from the National Research Foundation of Korea (NRF) grants funded by the Korea government (MSIT) (RS-2026-25515455). Joe Grove acknowledges support from the MRC-University of Glasgow Centre for Virus Research core support from the Medical Research Council (MC_UU_00034/1). Philippe Le Mercier acknowledges support by the Swiss State Secretariat for Education, Research and Innovation (SERI) and National Institute of Allergy and Infectious Diseases of the National Institutes of Health (U24AI183840). This work was supported by the BBSRC, UK Research and Innovation [20-BBSRC/NSF-BIO], Google DeepMind and the European Molecular Biology Laboratory. The AlphaFold Protein Structure Database is supported by Google DeepMind and the European Molecular Biology Laboratory.

## Competing interests

Risha Patel, Alexander I. Cowen-Rivers, Oleg Kovalevskiy, Agata Laydon, Jack Mason, Elisa L.H. Wong, and Jeremy Ratcliff are employees of Google DeepMind and may own stock as part of the standard compensation package. Riya Narain, Niccolò A. E. Venanzi, Darren Hsu, Nilkanth Patel, Kyle Gion, Davide Fransos and Christian Dallago are or were employees of NVIDIA. Martin Steinegger acknowledges outside interest in Stylus Medicine. The remaining authors declare no conflicts of interest.

## Methods

### Sequence and prediction candidate selection

There are 34 families of viruses known to infect humans (De Castro et al. 2024). To find well annotated sequence representatives across these families, we selected the International Committee on Taxonomy of Viruses (ICTV) Virus Metadata Resource (VMR; VMRv40.1) as a definitive and current record of exemplar species. The VMR also maps directly to the viral species included in UniProt, permitting us to derive proteome level sequence sets for each species. Many viruses encode simple open reading frames where one gene produces one protein, in these instances protein sequences can proceed directly to structure prediction. However, 15 of the human infecting families encode one or multiple polyproteins, where one gene produces a long polypeptide that is proteolytically cleaved to produce multiple mature proteins. In these cases accurate polyprotein sequence annotations are essential for biologically meaningful prediction of structure and interaction. The ViralZone team’s recent efforts have provided high-quality sequence annotations for four polyprotein encoding families, including the *Picornaviridae* which contains >500 exemplar species (**Supp. Fig. 2**). These well-annotated viruses were included in the current study, whilst the 11 families (∼700 species) without consistent and accurate annotations were excluded (e.g. *Coronaviridae* and *Retroviridae*). This resulted in a final input sequence set of 2,818 proteomes containing a total of 42,180 individual proteins, including 7,703 polyprotein mature products. Notably, VMRv40.1 contains the legacy-definition of the *Flaviviridae* prior to its recent taxonomic reorganisation (Simmonds et al. 2025). In this study we used the previous definition, in which the *Flaviviridae* contains four genera: hepacivirus, orthoflavivirus, pegivirus, and pestivirus. Monomer and homodimer candidates were derived from the 42,180 individual proteins. 1,677,356 heterodimer candidates were derived by pairing all proteins within a viral genome (intra-proteome all-vs-all). No inter-proteome heterodimers, including viral-host interactions were predicted.

### Benchmark dataset construction

Proximal pairs of chains from complexes of viral proteins were identified as a positive set by combining SIFTS-based PDB-to-UniProt mappings with structural proximity detected using Foldseek createdimerdb. Among 40,918 proximal chain pairs grouped into 5,870 structural clusters, the intersection of 19,780 viral PDB chain pairs mapped to UniProt via SIFTS yielded 17,619 candidate pairs supported by both sources. Structural clustering, redundancy reduction, representative selection, and post-training filtering against the PDB were performed following the previous approach (Humphreys et al. 2021; Han et al. 2026; Todor et al. 2026; J. Zhang et al. 2025). This yielded 2,798 representative PDB chain pairs, which were mapped to 943 unique pairs of UniProt accessions. After removing eight pairs containing host proteins, 935 viral pairs remained (571 bacteriophage, 361 non-phage, and three unclassified). From the 361 non-phage pairs, we retained non-polyprotein pairs with a BFVD v3 Logan-block match (R. S. Kim et al. 2026), yielding 215 positive pairs used in the final benchmark.

Negative pairs were derived from the same PDB structures supporting the 215 positive pairs. These positives mapped to 140 PDB structures containing 1,694 protein chains, from which 25,966 candidate chain pairs were enumerated. Pairs were excluded if they belonged to the same PDB entity, were spatially proximal according to createdimerdb, or the viral PDB interaction set mapped to UniProt via SIFTS, contained obsolete, sequence-unavailable, or manually excluded UniProt accessions, represented the same accession, had PDB-chain coverage < 70% of the corresponding UniProt sequence, or contained a protein ≥ 1,300 amino acids.

Potential near-identical sequence pairs were identified using MMseqs2 at sequence identity ≥ 0.9 and bidirectional coverage ≥ 0.8. Known interacting pairs were filtered using the full 935-pair viral set, the viral PDB interaction set mapped to UniProt via SIFTS, and createdimerdb pairs mapped to UniProt. This filter list was further expanded by substituting either or both proteins in each known interaction with their MMseqs2-defined sequence neighbors, increasing the filter list from 1,564 to 2,049 pairs. After all filters, 1,998 candidate negative pairs remained. Up to five negatives were selected per positive from its source PDB structure(s), prioritizing positives with smaller candidate pools and enforcing global accession-pair deduplication; random selection used seed 42. This yielded 242 unique negative pairs, from which pairs with a BFVD v3 Logan-block match were retained, resulting in 221 negative pairs used in the final benchmark. Negative labels indicate absence of spatial proximity in the observed PDB assemblies rather than absence of interaction in all biological contexts.

A pair was considered evaluable if prediction and all subsequent post-processing and metric evaluation steps were completed successfully. Of the 215 pairs in the positive set, 214 were evaluable for the analyses shown in **Fig. 2a,b**.

### Multiple Sequence Alignment generation

Multiple Sequence Alignments (MSAs) were generated for all 42,180 unique monomer sequences using colabfold_search (ColabFold 1.6.2) with the MMseqs2-GPU backend (Kallenborn et al. 2025), searching against UniRef30 and the ColabFold metagenomic database (--use-env 1). Dimer MSAs were generated directly in unpaired mode from concatenated two-chain input sequences. For polyprotein-encoding sequences, MSAs were generated directly from the mature product sequences (cleaved segments), as provided in the production FASTA, rather than from full-length polyprotein sequences followed by cropping.

### Structure prediction

Dimer structures were predicted using an accelerated implementation of OpenFold leveraging NVIDIA BioNeMo Inference Runtime with AlphaFold-Multimer v3 weights (model_1_multimer_v3) and 12 recycling iterations. Monomers were predicted using ColabFold v1.6.2 with alphafold2-ptm weights and 12 recycling iterations with early stopping. Failures due to out-of-memory errors or were observed in the presence of large MSAs and long sequences (greater than around 2100 amino acids). These failure modes occurred in 14,994 predictions and were not further investigated. Computation for any one complex was limited to four hours of execution time, and predictions exceeding this limit were not completed and excluded from downstream analysis. Among the 1,719,536 input dimeric pairs, a total of 1,704,542 dimeric structures were predicted, comprising 1,663,511 heterodimers and 41,031 homodimers.

### High-performance-compute scaling

MSA generation and structure prediction were scaled following (Han et al. 2026). Dimer predictions were distributed across NVIDIA DGX H100 and A100 nodes; pairs with combined sequence length ≤900 amino acids were predicted on A100 nodes, while longer pairs on H100 nodes.

### Post-processing toolkit and data products

A prior post-processing pipeline (Han et al. 2026) was extended to correctly propagate polyprotein fragment metadata (including sequence coordinates, protein name, and fragment status) through the chain-binding step, ensuring fragment segments were accurately represented in released data products. Models containing non-standard amino acids or coordinate-sequence mismatches were excluded prior to post-processing. Of 1,704,542 folded dimers, 1,699,311 (1,658,565 heterodimers and 40,746 homodimers) were successfully post-processed, of which 8,028 met the high-confidence threshold of ipSAE_max_ ≥ 0.6 and pDockQ2_max_ ≥ 0.23. Of 42,180 monomers, 41,869 were successfully post-processed. Among the 7,703 polyprotein monomeric entries, 146 predictions failed, leaving 7,557 successfully predicted monomeric structures. No quality thresholds were applied to monomers as ipSAE and pDockQ2 are interface metrics not applicable to single-chain structures.

### Assigning viral families

Every predicted model carries the NCBI taxonomy identifier (taxid) of the virus it came from, and the 1,699,311 models span 2,783 distinct identifiers. Each was resolved to a full lineage against a single fixed release of the NCBI taxonomy (downloaded 2026-09-09). Taxids that NCBI has since renumbered or withdrawn were followed through its merged- and deleted-node lists rather than discarded. 2,782 of the 2,783 taxids resolved to a family directly. The unresolved taxid had been withdrawn from the taxonomy and was assigned to the family shared by all other taxids from the same virus species. Viral identity was confirmed by tracing each lineage to the NCBI “Viruses” root node (taxid: 10239). This result yielded 25 families, which consolidated into 23 when nesting *Hepaciviridae* and *Pestiviridae* within *Flaviridae*, consistent with ICTV release VMR 40.1.

### Protein classification

Each modelled viral protein chain was identified by its UniProt accession and residue range, and classified as capsid, other or unclassified. Capsid-shell and core/nucleocapsid proteins were annotated separately but combined in the capsid category for compositional analyses. Non-polyprotein standalone proteins were classified using, in order of precedence, UniProt capsid keywords, capsid Gene Ontology (GO) cellular-component terms, nucleoprotein/nucleocapsid keywords and GO terms, and protein names. Annotations for capsid assembly or maturation, genome packaging, matrix, tegument and generic ribonucleoprotein complexes were not considered evidence of capsid identity. For name-based assignments, cleavage-product lists were removed from precursor names, and scaffolding and chaperone proteins were excluded from capsid assignment. Reviewed UniProtKB/Swiss-Prot entries lacking capsid or nucleocapsid evidence were assigned to the other category. Meanwhile, mature polyprotein products were only classified by product name rather than precursor-level annotations, using a classification table covering all 77 distinct product names in the dataset (**Supp. Table 1**). Chains without an assignment, including polyprotein chains spanning multiple mature products, remained unclassified. The supporting evidence was recorded for each chain. Among 41,774 distinct chains from 34,964 UniProt accessions represented across the 1,699,311 predicted dimers, 7,183 were classified as capsid proteins (6,606 capsid and 577 core/nucleocapsid), 31,610 as other, and 2,981 as unclassified. Dimers were classified as capsid–capsid, capsid–other or other–other; dimers containing an unclassified chain were reported separately.

### Interface Clustering of PDB

PDB structures downloaded in January 2026 were processed using foldseek (commit d6204679), following the pipeline and parameters described in the Foldseek-Interface ((Strom et al. 2026)). Dimers extracted using createdimerdb were clustered with easy-multimercluster at TM-score and chain TM-score thresholds of 0.9 and 0.5, respectively. Cluster representatives were compiled into repdb, and interfaces were extracted using createinterfacedb. These interfaces were then clustered with easy-multimercluster using exhaustive search, TM-score and lDDT thresholds of 0.4 and 0.20, respectively, and an E-value cutoff of 10⁷, yielding 118,269 interface clusters.

### Interface clustering and search of the high-confidence subset

The 8,028 high-confidence viral dimers (ipSAE_max_ ≥ 0.6; pDockQ2_max_ ≥ 0.23) were clustered using Foldseek (commit d6204679), following the two-stage pipeline and parameters described by Strom et al. ((Strom et al. 2026)). First, the di-mer clustering step yielded 4,951 clusters (a 1.6-fold reduction), followed by an interface clustering step that yielded 1,598 interface clusters.

Interfaces extracted from all 8,028 high-confidence viral dimers were searched against 118,269 PDB interface cluster representatives. This search followed the HumanPPI-to-PDB procedure described by Strom et al. (Strom et al. 2026), using easy-multimersearch (Kim et al., 2025) with --cov-mode 0 and otherwise default search parameters. A PDB hit was defined as max(qTM, tTM) > 0.4, where qTM and tTM denote query- and target-normalised TM-scores, respectively. A cluster was considered PDB-matched if at least one member had a qualifying hit. Overall, 6,598 dimers (82.2%) had PDB hits, whereas 471 of the 1,598 interface clusters (29.5%) had no PDB hit (**Fig. 4a**).

### AdV protease/substrate alignment and sequence logo analysis

Substrate sequences were aligned by structure, rather than by sequence, using a single large interface-cluster (n=69) of AdV protease/substrate pairs. For each complex we measured the distance from the catalytic cysteine sulfur to the backbone carbonyl carbon of every substrate residue and took the nearest as an anchor, this being the carbon a cysteine protease attacks. Each substrate sequence was then read as a window of three residues either side of the anchor, and the windows stacked, so every column holds the same offset from the anchor. Because the anchor is measured in each structure independently the resulting sequence motif arises purely from geometric arrangement in the active site. Of the 69 models in the cluster, 65 were used; the remaining four were excluded because no part of the substrate reaches the active site. Per column, Shannon information content was calculated with the standard small-sample correction, and logos were drawn with logomaker (Tareen and Kinney 2020), with letters scaled by frequency and coloured by residue class.

### Molecular modelling

Visualisation of protein structures was performed with Mol* (Sehnal et al. 2021).

### Supplementary Figures

**Supplementary Figure 1.**
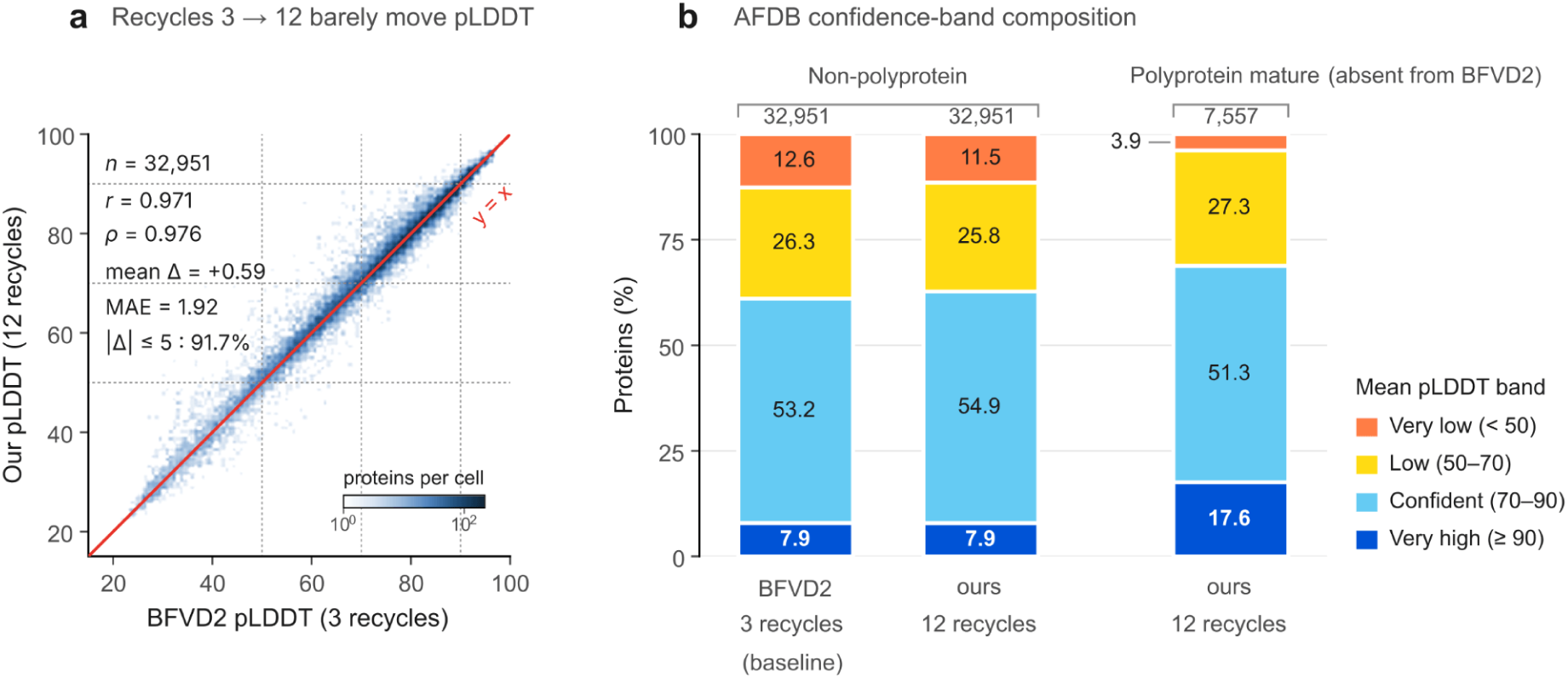
Monomer confidence is robust to recycling depth and elevated for mature polyprotein products. **a**, Of the 41,869 post-processed monomeric predicted structures,32,957 were mappable to BFVD v3 ((R. S. Kim et al. 2025; Litvin et al. 2025; Odai et al. 2025; R. S. Kim et al. 2026)). Mean pLDDT for 32,951 non-polyprotein viral proteins predicted with three recycles (BFVD v3) or 12 recycles (used in this study), with otherwise identical ColabFold settings. Predictions were highly concordant (Pearson *r* = 0.9708; Spearman ρ = 0.9756; mean absolute difference, 1.92 pLDDT), with a mean increase of 0.59 pLDDT at 12 recycles; 91.72% differed by ≤5 pLDDT and 91.67% remained in the same AlphaFold DB confidence category. Density is shown on a logarithmic scale; the red line denotes *y* = *x* and dashed lines mark pLDDT thresholds of 50, 70 and 90. **b**, Distribution across AlphaFold Database confidence categories. Increasing from three to 12 recycles reduced the very-low-confidence fraction by 1.2 percentage points but left the very-high-confidence fraction unchanged (7.89% versus 7.88%). Mature polyprotein products (*n* = 7,557) were more often very high confidence (17.6% versus 7.9%) and less often very low confidence (3.9% versus 11.5%) than non-polyprotein proteins. BFVD v3 contains no equivalent mature-product set. Numbers denote protein counts and percentages. pLDDT measures model confidence, not structural accuracy.

**Supplementary Figure 2.**
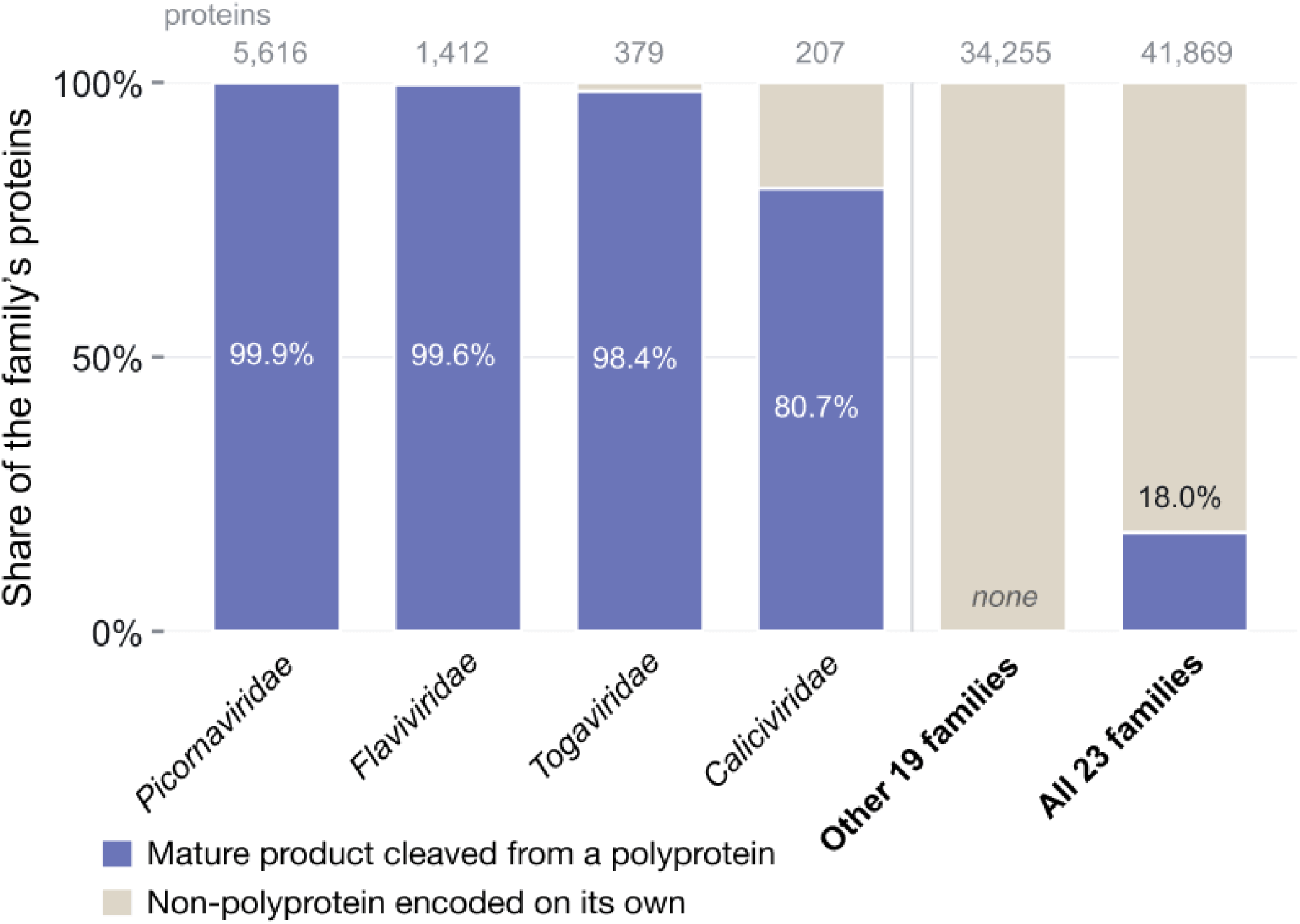
Polyprotein content of each viral family. Four of 23 viral families encode polyproteins, which accounts for 81-100% of their proteins. In contrast, the other 19 families, holding 34,160 proteins, do not encode polyproteins. Of the 41,869 post-processed monomeric predicted structures, 7,557 (18.1%) corresponded to polyprotein mature products. Families follow VMR 40, which places *Hepaciviridae* and *Pestiviridae* inside *Flaviviridae*; this merge reduces the number of families with polyproteins from 6 to 4.

**Supplementary Figure 3.**
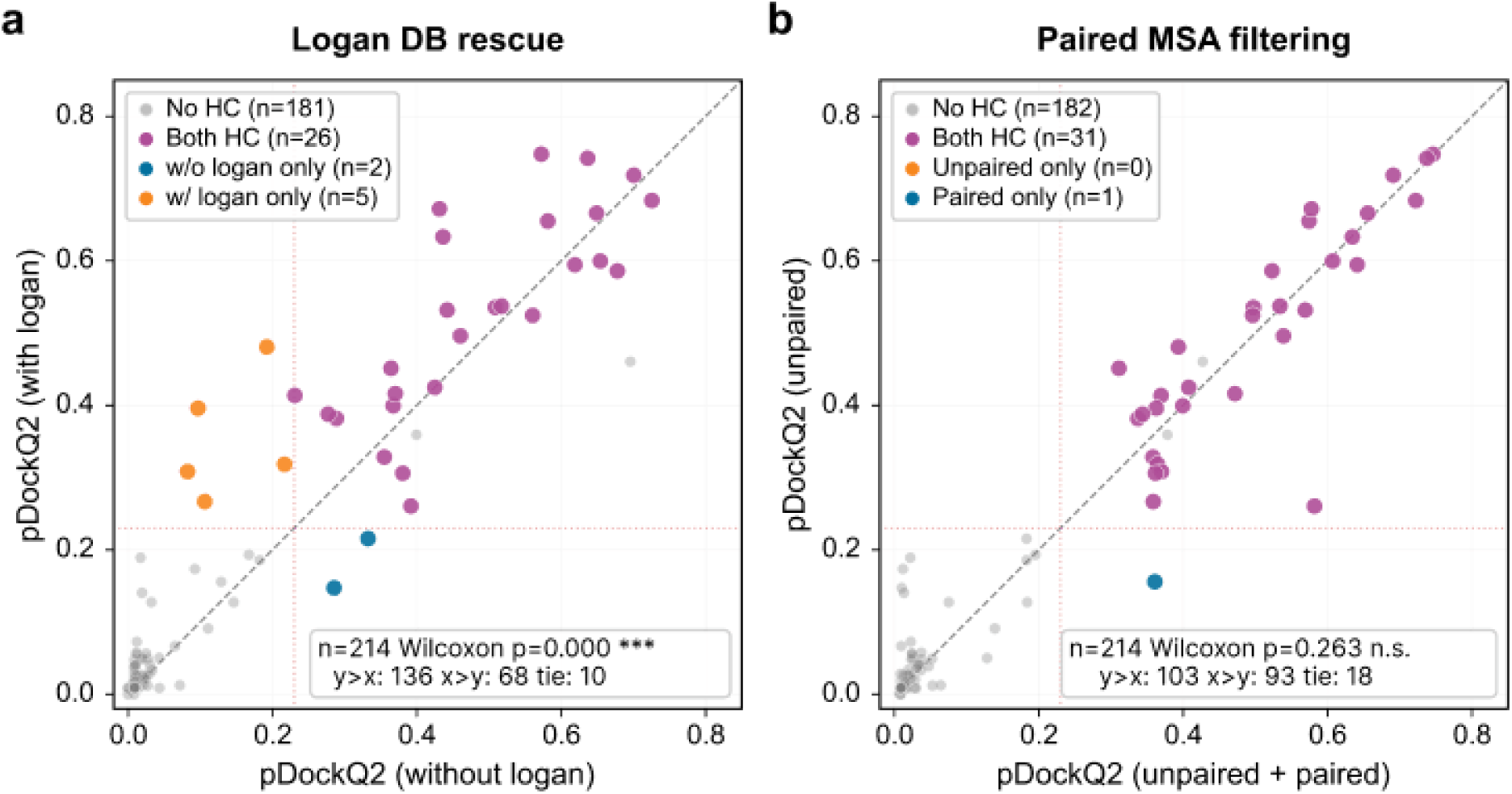
pDockQ2-based evaluation of MSA curation strategies. Same analysis as **Fig. 2a,b**, displaying pDockQ2 rather than ipSAE. High-confidence classification remains defined jointly by ipSAE ≥ 0.6 and pDockQ2 ≥ 0.23; dotted lines mark the pDockQ2 threshold. **a,** Effect of LoganDB enrichment. pDockQ2 was significantly higher with LoganDB enrichment (Wilcoxon signed-rank test, *P* < 0.001), increasing in 136 of 214 benchmark pairs. High-confidence classifications were the same to **Fig. 2a** (both, *n* = 26; ‘with Logan’ only, *n* = 5; ‘without LoganDB’ only, *n* = 2). **b,** Effect of paired MSA filtering. pDockQ2 scores were similar between filtered paired-plus-unpaired and unpaired-only MSAs (Wilcoxon signed-rank test, *P* = 0.299), consistent with the ipSAE analysis in **Fig. 2b**.

**Supplementary Figure 4.**
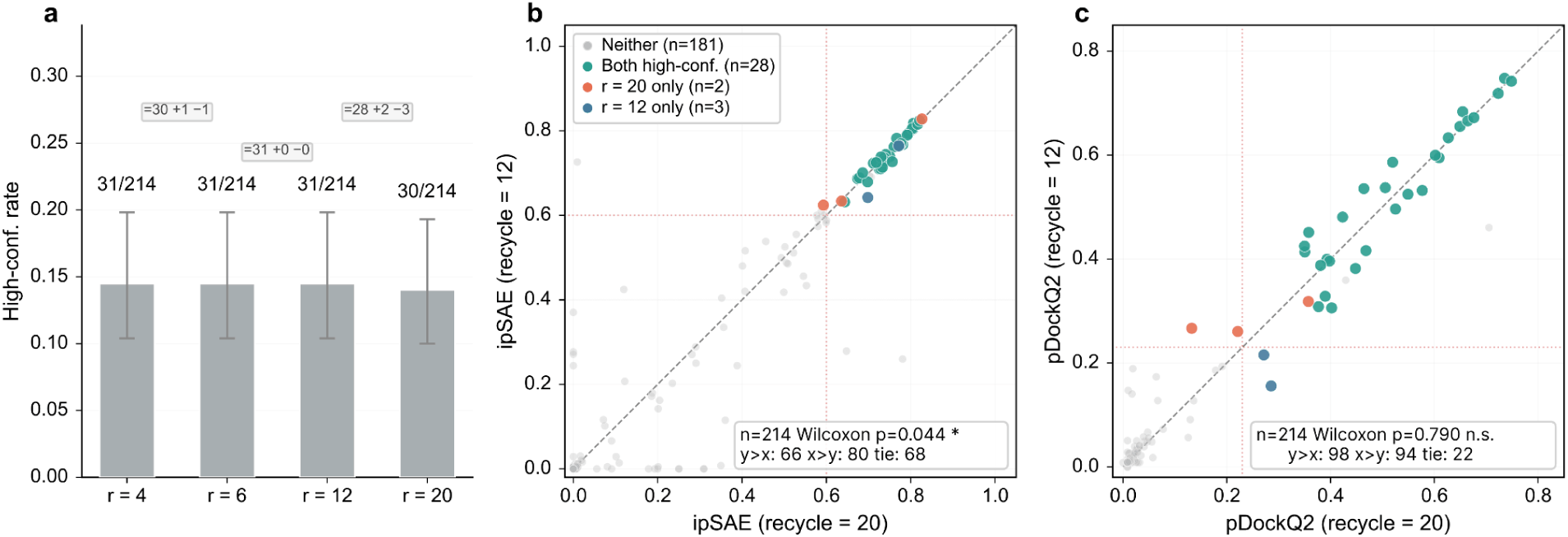
High-confidence classification is robust to the number of recycling iterations. **a**, Fraction of high-confidence predictions across 4, 6, 12 and 20 recycling iterations for the unpaired-MSA pipeline (*n* = 214 evaluable non-phage benchmark pairs). High confidence was defined jointly as ipSAE ≥ 0.6 and pDockQ2 ≥ 0.23. Error bars indicate 95% Wilson confidence intervals. Annotations show the numbers of concordant (=), gained (+) and lost (−) high-confidence pairs between consecutive settings. Counts were nearly invariant (31, 31, 31 and 30), with complete concordance between 6 and 12 recycles. **b,c,** Per-pair comparisons of ipSAE (b) and pDockQ2 (c) at 20 recycles (x axis; highest recycling depth tested) and 12 recycles (y axis; setting used in this study). Points are coloured by joint classification: teal, high confidence at both settings; coral, 20 recycles only; steel blue, 12 recycles only; grey, neither.

**Supplementary Figure 5.**
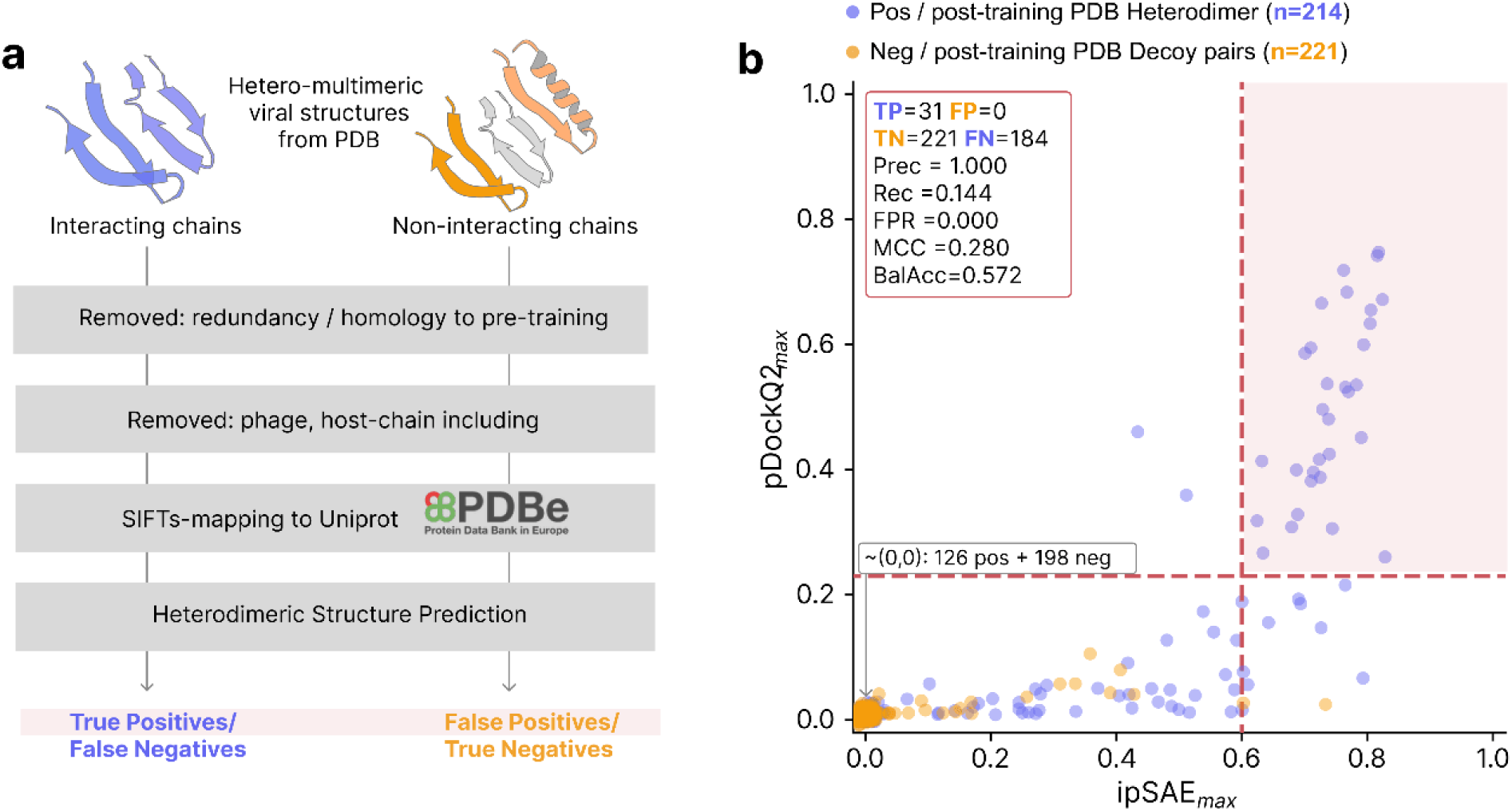
Post-training benchmark of the viral dimer prediction pipeline. **a**, Benchmark workflow. Positive (n = 215) and negative (n = 221) pairs were derived from post-training PDB structures, mapped to UniProt using SIFTS and modelled with our prediction pipeline. Structures similar to pre-training complexes were excluded using Foldseek-Multimer searches (Methods). Classification used 215 positive and 221 evaluable negative pairs. **b,** Joint distribution of ipSAE_max_ and pDockQ2_max_ for positive (purple) and negative (orange) pairs. Each metric is the maximum across the two chain directions. Dashed lines mark the joint high-confidence criterion (ipSAE_max_ ≥ 0.60 and pDockQ2_max_ ≥ 0.23). High-confidence positives and negatives are true positives (TP = 31) and false positives (FP = 0), respectively; the remaining evaluable pairs are false negatives (FN = 184) and true negatives (TN = 221). Points near the origin are jittered for visibility; classification uses unmodified scores.

**Supp. Fig 6.**
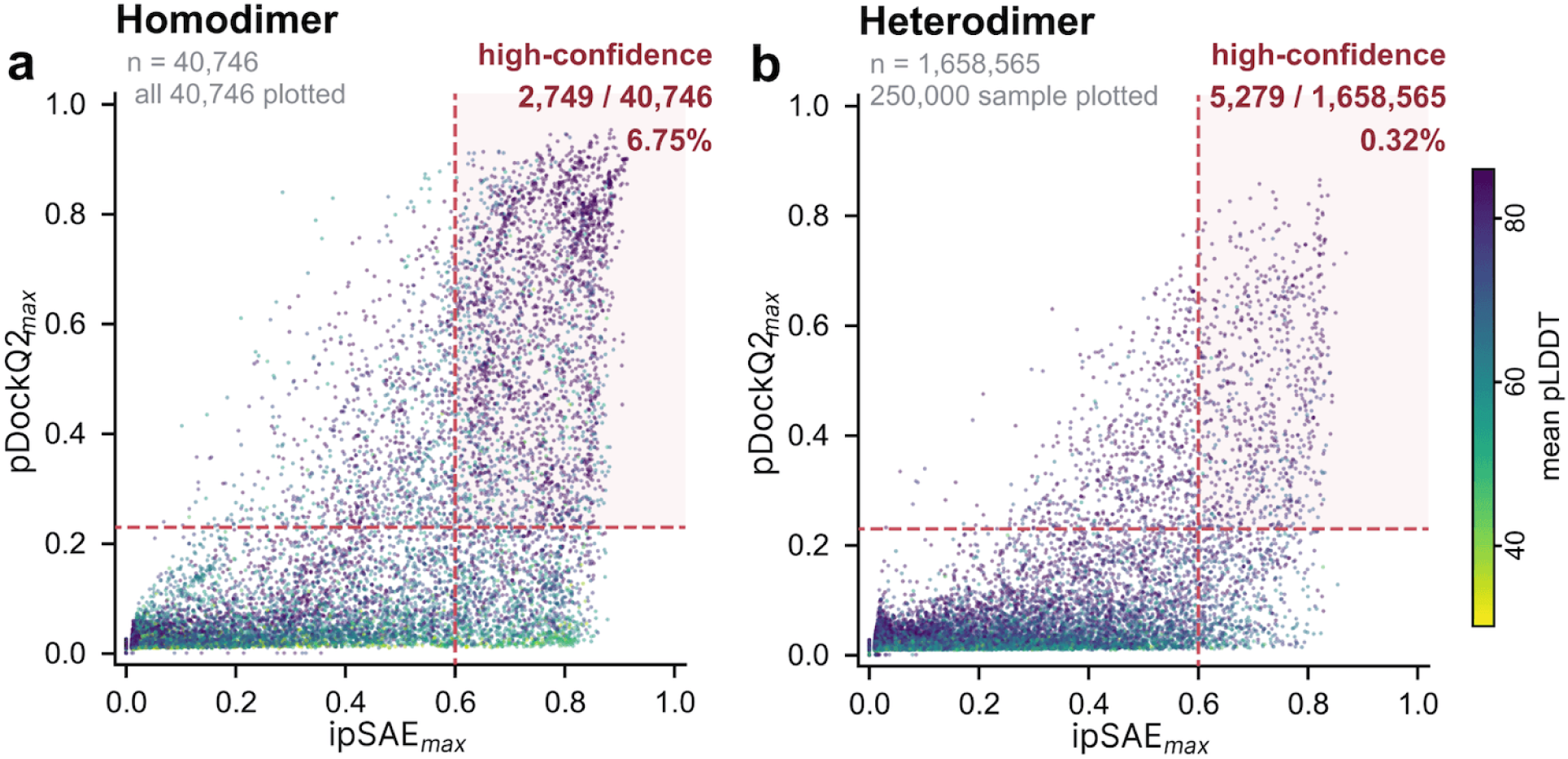
Homodimer & heterodimer production result. Predicted interface scores for every homodimer (a) and heterodimer (b) model, coloured by mean pLDDT. Dashed lines mark the two thresholds; the upper-right quadrant is the high-confidence set, with its count and rate given in place. Points are a random sample of at most 250,000 models per panel; all counts and rates are computed on the full population, not the sample.

**Supplementary Figure 7.**
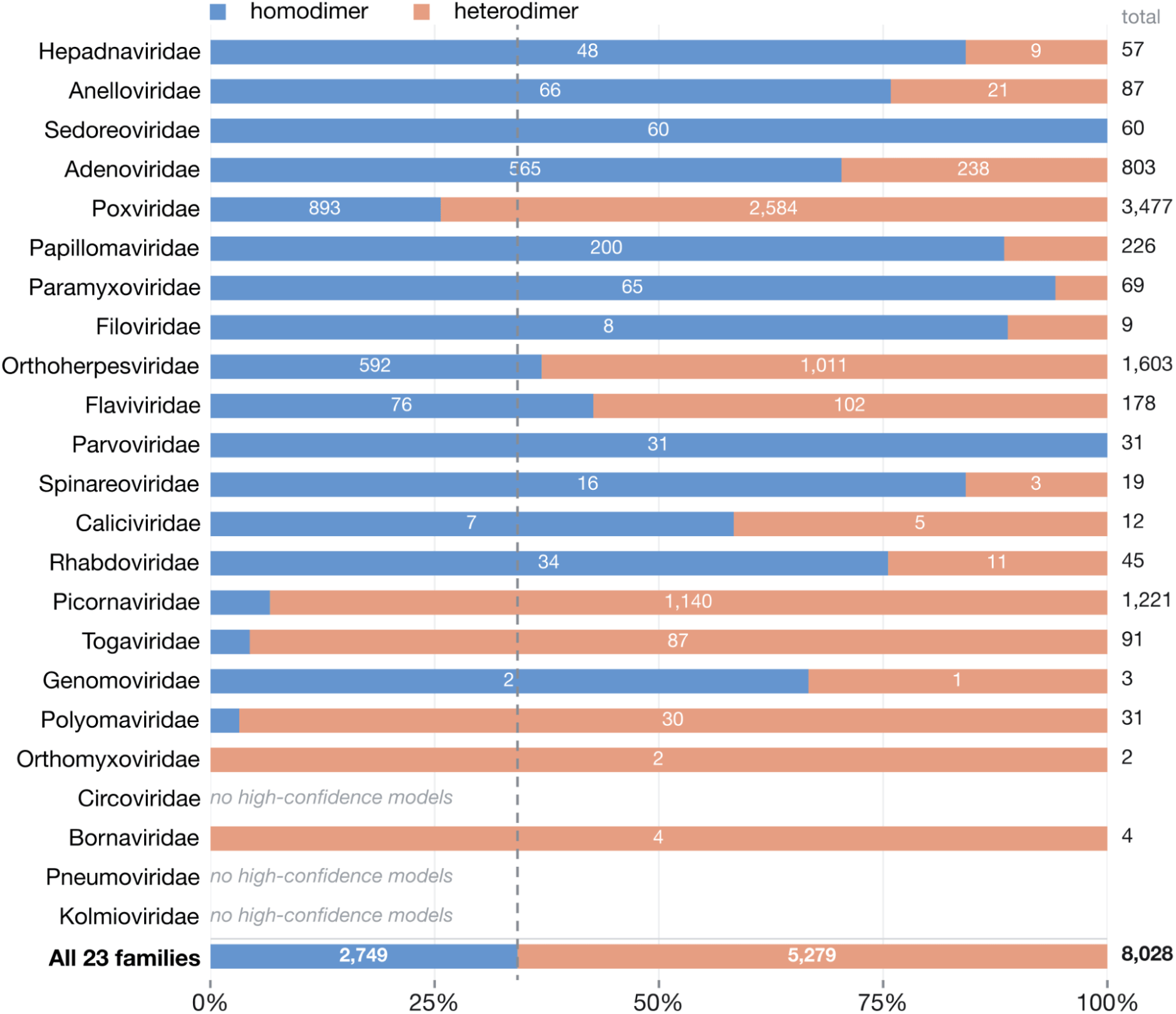
Composition of the high-confidence predictions in each viral family. Each bar shows how one family’s high-confidence models divide between pairs of a protein with a copy of itself and pairs of two different proteins of the same virus, scaled to 100%; the two counts are given inside the bar and the family total at the right. The dashed line marks all 23 families combined, shown in the bottom row. 3 families yielded no high-confidence model and so have no composition. Families are ordered as in **Fig. 3a**. Pairs of identical proteins make up 2.4% of the 1,699,311 predictions but 34% of the 8,028 high-confidence ones. The proportion differs widely between families and does not follow the ranking by success rate: *Poxviridae* contributes the most high-confidence models (3,477), of which 26% are identical pairs, while *Hepadnaviridae*, which has the highest success rate for identical pairs in **Fig. 3a**, contributes 57. Numbers of predictions attempted and success rates for every family are given in the accompanying table.

**Supplementary Figure 8.**
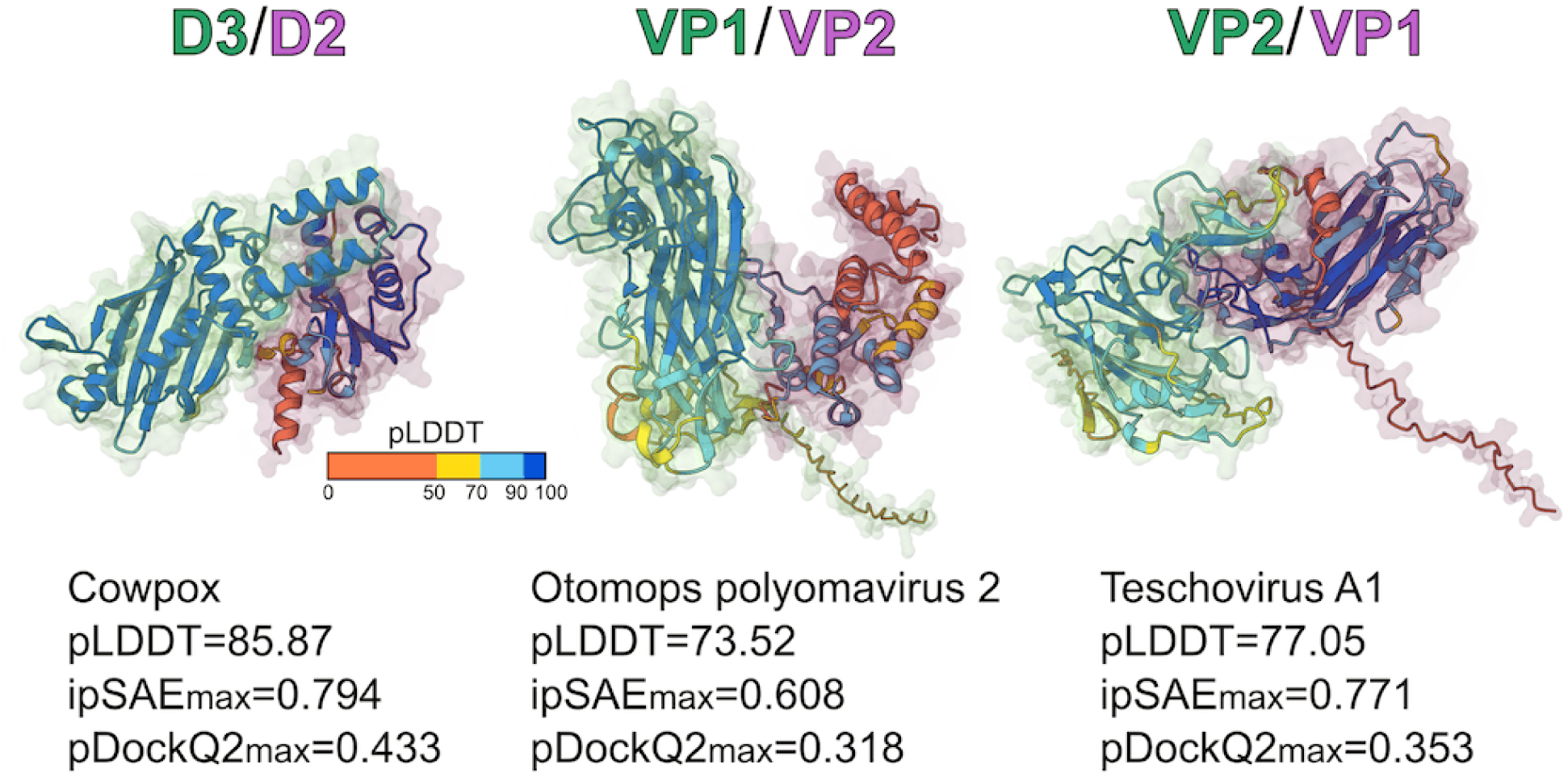
Capsid assemblies. Example high-confidence viral capsid protein heterodimers, from diverse viruses, that are otherwise absent or poorly represented on the PDB (from left to right: AF-0000000213303165, AF-0000000212018239, ,AF-0000000212074116). Ribbon diagrams are colour-coded by pLDDT prediction confidence, as denoted in the key, with transparent surface representations highlighting individual chains colour-coded by protein identity, as shown in the complex labels. Annotations provide average pLDDT values and interaction confidence metrics.

**Supplementary Figure 9.**
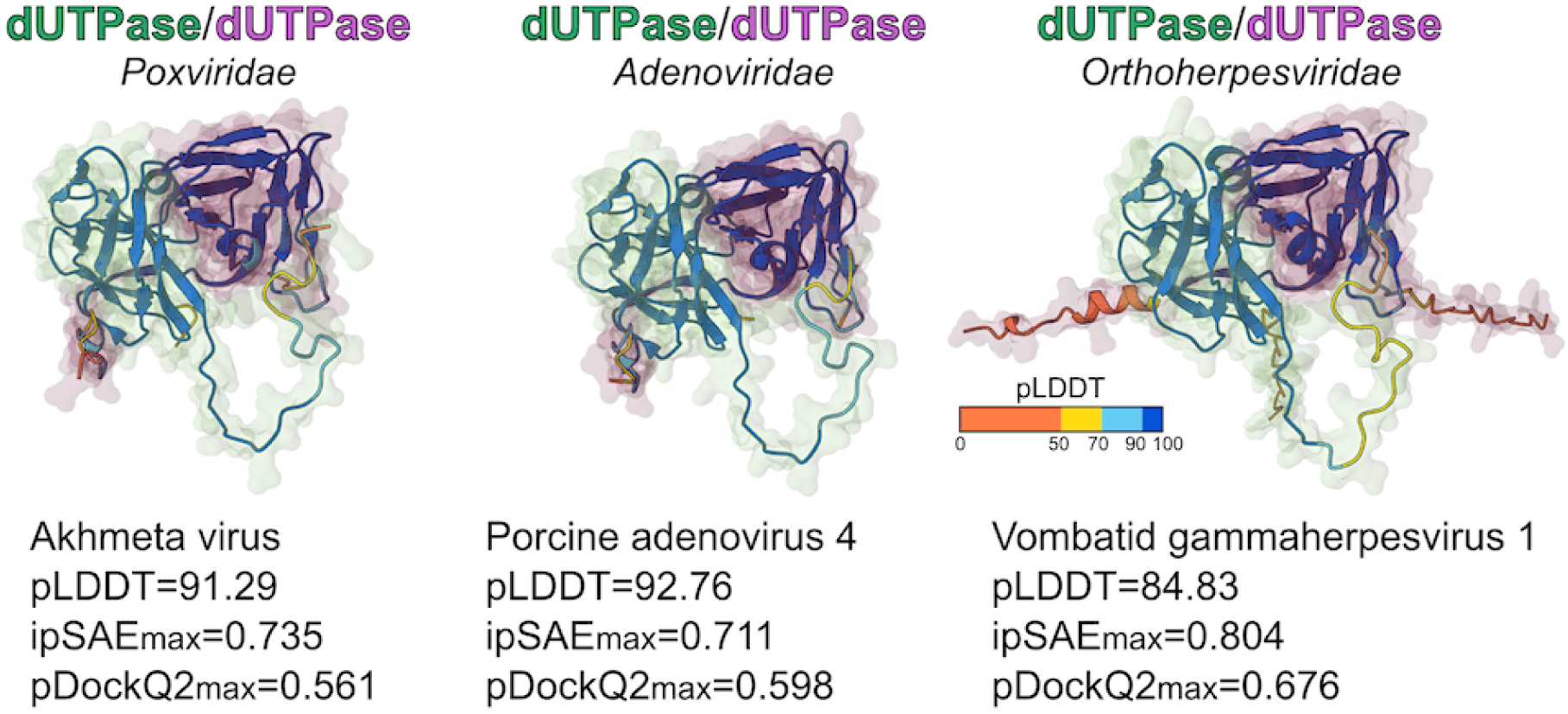
A cross-family interface-cluster of viral dUTPase homodimers. Example structures from an interface-cluster of dUTPase homodimers, containing members from the *Poxviridae* (AF-0000000213202665), *Adenoviridae* (AF-0000000212119938), *Orthoherpesviridae* (AF-0000000212243866). Ribbon diagrams are colour-coded by pLDDT prediction confidence, as denoted in the key, with transparent surface representations highlighting individual chains colour-coded by protein identity, as shown in the complex labels. Annotations provide average pLDDT values and interaction confidence metrics.

**Supplementary Figure 10.**
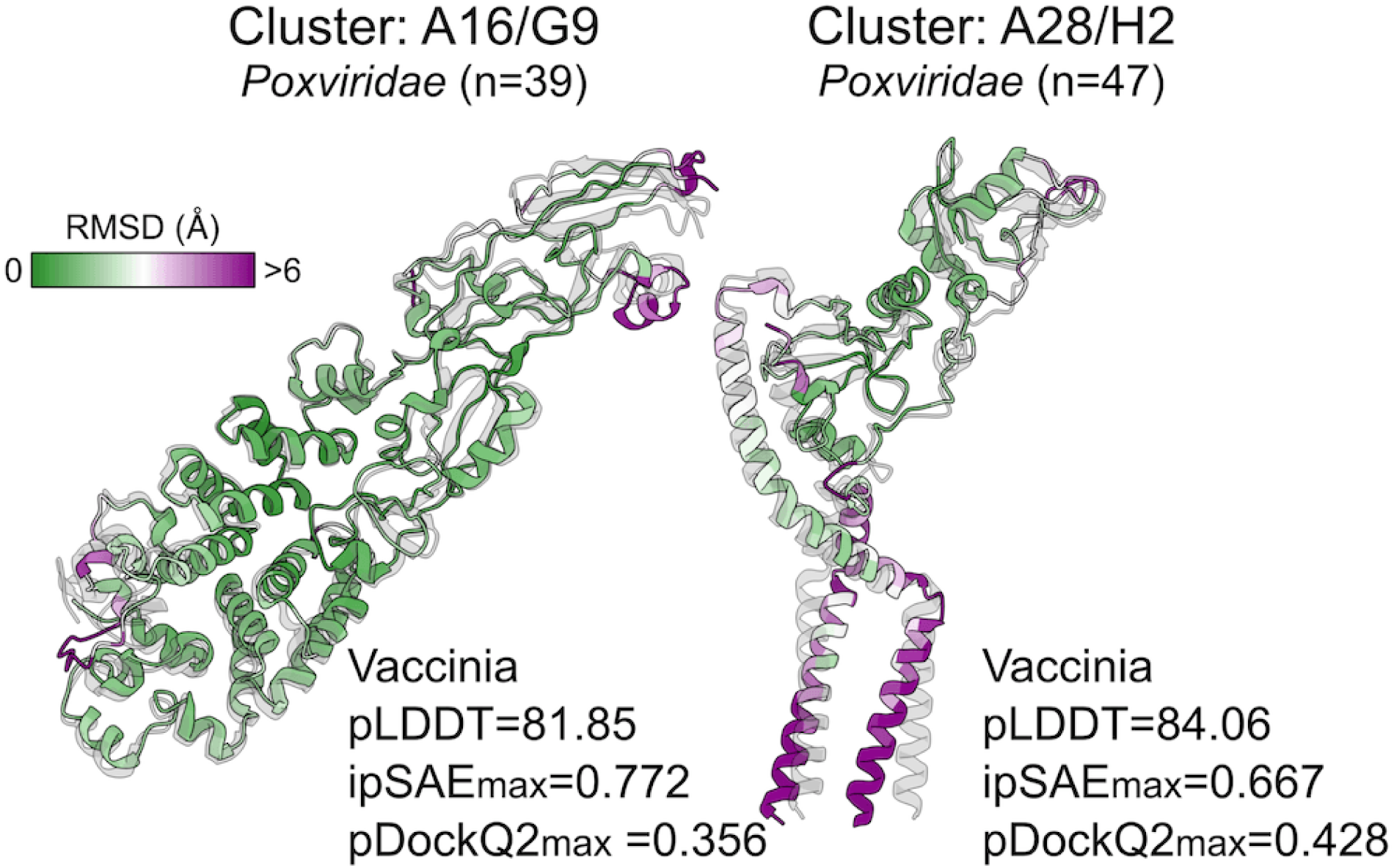
Comparison of vaccinia EFC predictions with recent experimental structures. Dimer structures taken from two interface clusters representing components of the poxvirus entry fusion complex, A16/G9 and A28/H2, as in **Fig. 4b**. Ribbon diagrams are colour coded by C-⍺ RMSD following superposition with recently published experimental structures, PDB references are shown as grey transparent ribbons (A16/G9 vs pdb_00009R0J, A28/H2 vs pdb_00009UZP). Regions of heterodimers that are not present in the experimental structures have been excluded from the visualisation. Annotations provide average pLDDT values and interaction confidence metrics. Cluster labels indicate complex identity, viral taxonomy and, in parentheses, number of members per cluster.

**Supplementary Table 1.** Classification of polyprotein mature-product names. All 77 distinct mature-product names in the dataset, grouped by the category assigned to them and together covering 7,557 chains. Products of a polyprotein are classified from the product name, because a UniProt keyword or Gene Ontology term attaches to the whole polyprotein and cannot indicate which product a chain is. Capsid and core are reported separately here and combined for compositional analyses; unclassified means the name does not identify the product. Criteria for proteins that are not polyprotein products are given in the accompanying table.

| CATEGORY | PRODUCT NAMES | NAMES | CHAINS |
| --- | --- | --- | --- |
| Capsid | Capsid protein, Capsid protein C, Capsid protein VP0, Capsid protein VP1, Capsid protein VP2, Capsid protein VP3, Capsid protein VP4, Capsid protein VP60 | 8 | 2,028 |
| Core / nucleocapsid | Core protein, Mature core protein | 2 | 26 |
| Other | 3C like protease, 6K protein, Assembly protein E3, Assembly signal 2A, Cysteine protease NS2, E rns glycoprotein, Envelope glycoprotein E1, Envelope glycoprotein E2, Envelope protein E, Leader protease, Leader protein, N terminal protease, NS1, NS1 2, NS1 NS2, NS2, NS4, NTPase, Non structural protein 1, Non structural protein 2A, Non structural protein 3, Non structural protein 4A, Non structural protein 4B, Non structural protein 5A, Peptide 2k, Peptide pr, Protease 2A, Protease 3C, Protease NS2, Protease nsP2, Protease polymerase, Protein 2A, Protein 2A 1, Protein 2A 2, Protein 2A 3, Protein 2A 4, Protein 2A H NC, Protein 2A NTPase, Protein 2B, Protein 2C, Protein 3A, Protein 3B, Protein 3B 1, Protein 3B 2, Protein 3B 3, Protein PrM, Protein prM, RNA directed RNA polymerase, RNA directed RNA polymerase 3D POL, RNA directed RNA polymerase Methyltransferase NS5, RNA directed RNA polymerase nsP4, Serine protease Helicase NS3, Serine protease NS3, Serine protease helicase NS3, Serine protease subunit NS2B, Small envelope protein M, Spike glycoprotein E1, Spike glycoprotein E2, VPg, Viral genome linked protein, Viral protein genome linked, Viroporin p7, Viroporin p7 like, mRNA capping enzyme nsP1, p13, p6 | 66 | 5,483 |
| Unclassified | Protein Y | 1 | 20 |

## Notes

https://alphafold.ebi.ac.uk/search/thematic-tag/pandemic-preparedness

